# Evolution of an Aurora Kinase A Inhibitor from an Essential tRNA Synthetase

**DOI:** 10.64898/2026.05.15.725417

**Authors:** James Julian Ross, Polina Tikanova, Andreas Hagmüller, Daniel Krogull, Keyu Xiao, Casper van Bavel, Till Balla, Tianhua Liao, Kolin Angelo Echano, Joseph Gokcezade, Peter Duchek, Danny Nedialkova, Rob Jelier, Gang Dong, Alejandro Burga

## Abstract

Gene duplication is a major driver of evolution, yet how it generates fundamentally new molecular functions remains poorly understood. Here, we show how such novelty arose in KLMT-1, a selfish toxin that causes genetic incompatibilities in *Caenorhabditis tropicalis*. KLMT-1 evolved via duplication of an essential tRNA synthetase but, strikingly, lost its ancestral role in tRNA biology and translation. Instead, KLMT-1 localizes to centrosomes, where it targets Aurora kinase A (AIR-1). This innovation is mediated by a three–amino acid insertion that extends a β-hairpin loop, enabling electrostatic interaction with a regulatory interface on the kinase. Our results demonstrate how changes in selective pressure, combined with minimal modifications in neutrally evolving regions, allow duplicated proteins to access new functional space and evolve entirely new molecular activities.

## Main text

Born equal and liberated from the restraints of purifying selection, duplicated genes are free to diverge and explore new functions. For this reason, gene duplication is regarded as one the major forces driving evolutionary innovation (*1–4*). Yet the apple rarely falls far from the tree. Duplicated transcription factors may bind new DNA motifs, receptors may recognize novel ligands, enzymes may shift their substrate selectivity—changes that are important, but still variations on the same ancestral theme. This raises a critical question: if gene duplication largely modifies what already exists, to what extent can it give rise to functions that are genuinely new?

At the molecular level, most well-documented cases of novelty following gene duplication build on pre-existing functions. Innovation usually arises through the partitioning of ancestral protein domains between paralogs or through the acquisition of new domains by recombination with other genes (*5–10*). Another recurring pattern is that ancestral proteins harbour secondary activities that can be amplified after duplication. This allows one copy to specialize on the secondary function while the other retains the original role, thereby resolving an adaptive conflict. The few known cases include the evolution of antifreeze proteins in fish, lens crystallins in vertebrates, and biosynthetic enzymes in plants (*11–13*). Despite these insights, how gene duplication leads to radically new molecular functions remains poorly understood, in large part because clear-cut examples are exceedingly rare (*14*, *15*). Here, we uncover at the molecular level one such critical event, in which a recent duplication of an enzyme essential for translation gave rise, through evolution of a novel binding interface, to a toxic inhibitor of Aurora kinase A in centrosomes.

### The KLMT-1 toxin evolved from FARS-3 by gene duplication

Toxin–antidote (TA) elements are selfish genes that subvert the laws of Mendelian inheritance and, in doing so, cause genetic incompatibilities in a wide range of species (*16*, *17*). While studying hybrids between Caribbean isolates of the nematode *Caenorhabditis tropicalis*, we recently identified one such element, the *klmt-1/kss-1* gene pair (Fig. 1A) (*18*, *19*). *klmt-1* encodes a maternally expressed toxin, whereas *kss-1* encodes its zygotically expressed antidote. In crosses between carrier and non-carrier strains, heterozygous F_1_ mothers deposit KLMT-1 into all eggs, effectively poisoning every embryo. Because the toxin and antidote genes are genetically linked, only progeny that inherit the TA locus can express KSS-1 and survive, whereas virtually all homozygous non-carriers—25% of the F_2_ generation—lack the antidote and die, leading to preferential transmission of the selfish locus (Fig. 1A,B).

**Fig. 1.**
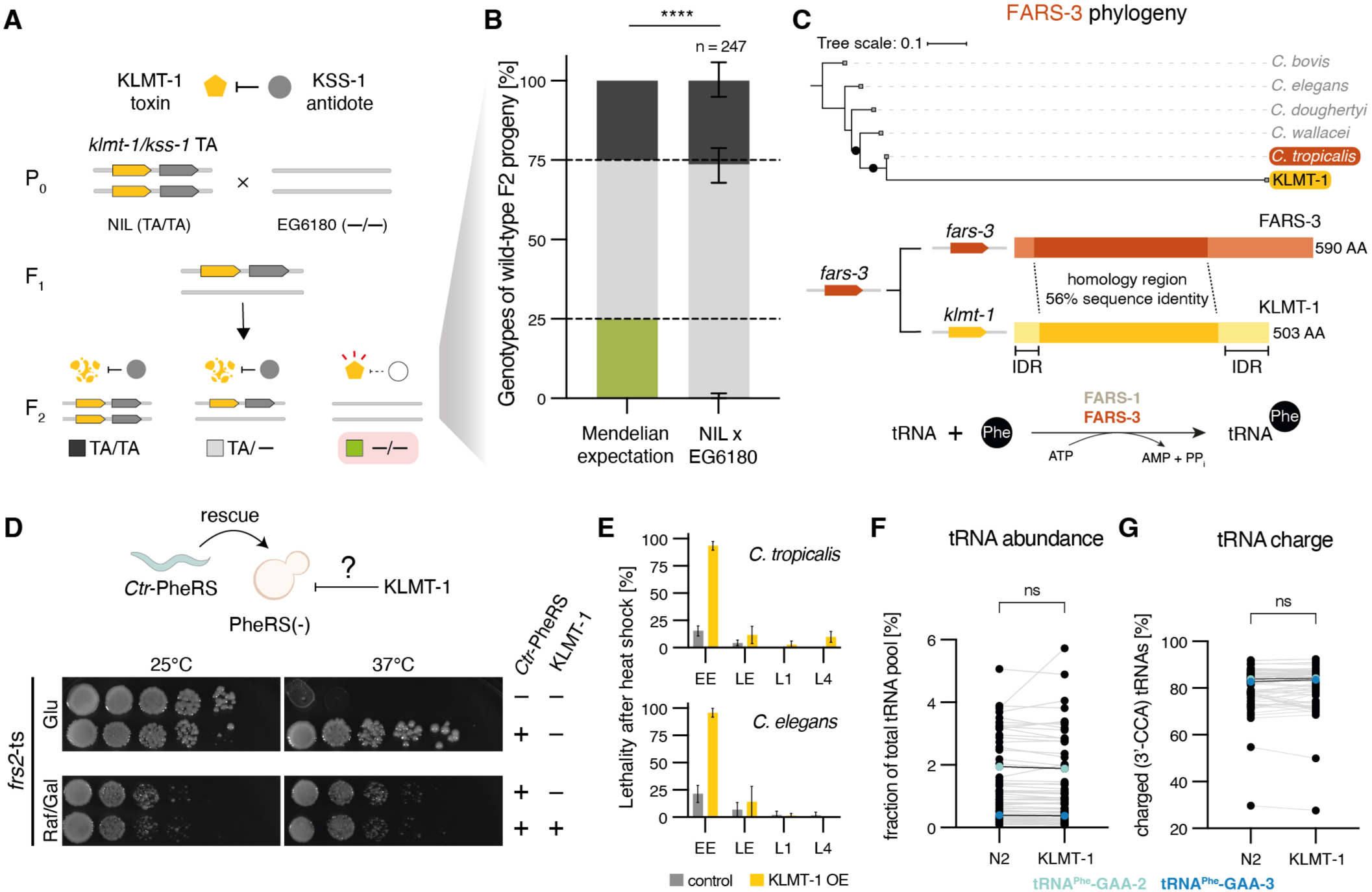
The KLMT-1 toxin evolved from a tRNA synthetase but it does not hinder the function of tRNAs. **(A)** Schematic of a toxin-antidote (TA) element. EG6180 worms containing a *klmt-1/kss-1*-containing introgression from NIC203 on Chr. IV (NIL) are crossed to the susceptible line EG6180 (EG). All F2 progeny are exposed to the maternally deposited toxin, yet only EG/EG individuals are affected since they cannot zygotically express the antidote. **(B)** The *klmt-1/kss-1* TA system acts as a highly efficient Mendelian distorter. In crosses between a *klmt-1/kss-1* carrier and susceptible parental strains, F_2_ individuals homozygous for the susceptible allele (−/−) die as embryos or L1 larvae (*n* = 247, *P* < 0.0001, chi-square test). Error bars represent the 95% confidence interval (hybrid Wilson/Brown). For this and all following figures: \**P* < 0.05, \*\**P* < 0.01, \*\*\**P* < 0.001, \*\*\*\**P* < 0.0001, ns = not significant. **(C)** Phylogenetic tree of KLMT-1 and FARS-3 (top). Comparison of protein coding regions of KLMT-1 and *C.tr*-FARS-3. Intrinsically disordered region (IDR) (middle). Simplified schematic of the tRNA charging reaction catalyzed by the PheRS (bottom). **(D)** Co-expression of *C. tropicalis* FARS-1 and FARS-3 rescues growth of *S. cerevisiae frs2*(ts) mutant at the restrictive temperature (37°C) in glucose-containing medium (top). Additional expression of KLMT-1 under an inducible galactose promoter did not affect growth of the rescued strain in raffinose/galactose (raf/gal) medium (bottom) **(E)**. Quantification of lethality upon heat shock Induction of a single copy *hsp-16.2*p::*klmt-1* transgene in different developmental stages of *C. tropicalis* (top) and *C. elegans* (bottom). Data from three individual experiments. EE = early embryos, LE = late embryos, L1/L4 = larval stages. Control is heat shock of WT parental line. **(F)** Abundance of all major *C. elegans* tRNA species (>0.1% of total pool) measured by mim-tRNAseq in heat-shocked wild-type embryos (control) or heat-schocked *hsp-16.2*p::*klmt-1* transgenic embryos, each in biological quadruplicates. KLMT-1 does not affect tRNA abundance (*P* = 0.974, paired t-test). tRNA^Phe^ isodecoders are highlighted in shades of blue. **(G)** Fraction of charged molecules for each *C. elegans* tRNA species measured by mim-tRNAseq in heat-shocked wild-type embryos (control) or heat-schocked *hsp-16.2*p::*klmt-1* transgenic embryos, each in biological quadruplicates. KLMT-1 does not affect charge ratios (P = 0.514, paired t-test). Highlight colours as in F.

The antidote, KSS-1, is a rapidly evolving F-box protein that binds KLMT-1 and targets it for proteasomal degradation by recruiting the SCF ubiquitin ligase complex (*18*). On the other hand, *klmt-1* arose via gene duplication from a highly conserved gene, *fars-3*, an essential subunit of the phenylalanyl-tRNA synthetase (PheRS) complex (Fig. 1C). PheRS plays an essential role in mRNA translation by charging tRNA^Phe^ with its cognate amino acid, L-phenylalanine (Fig. 1C). In bacteria, archaea and eukaryotes, PheRS is a heterotetrameric enzyme made up of two α (FARSA) and two β (FARSB) subunits, which in nematodes are encoded by *fars-1* and *fars-3,* respectively (*20*, *21*). FARSB is essential for tRNA^Phe^ aminoacylation through its obligate interface with the catalytic FARSA subunit, which stabilizes the active enzyme complex. This interface constrains tRNA^Phe^ positioning for efficient phenylalanine transfer and enables proofreading of mis-aminoacylated tRNA species (*22*). Based on their evolutionary relationship and shared structural features (Fig. 1C) (*18*), we hypothesized that KLMT-1 exerts its toxicity by competing with FARS-3 for binding to FARS-1, thereby interfering with PheRS assembly and acting as a dominant-negative form.

### KLMT-1 does not disrupt tRNA^Phe^ charging or tRNA function

To test whether KLMT-1 inhibits the *C. tropicalis* PheRS, we developed a heterologous complementation assay in *S. cerevisiae*. We reasoned that if the yeast PheRS were functionally replaced by its *C. tropicalis* counterpart, KLMT-1 should impair yeast growth if the PheRS is its direct target. To implement this, we used a yeast strain carrying a temperature-sensitive allele of the PheRS α-subunit (*frs2-ts*), which grows at 30°C but exhibits lethality at 35°C (*23*). Expression of *C. tropicalis* FARS-1 and FARS-3 from a plasmid fully rescued growth of these mutants at the restrictive temperature, indicating that the *C. tropicalis* PheRS can efficiently charge yeast tRNA^Phe^ (Fig. 1D). We then introduced a second plasmid expressing KLMT-1 under a GAL-inducible promoter and compared yeast growth with or without the toxin. Although *C. tropicalis* PheRS activity was essential for yeast viability at the restrictive temperature and KLMT-1 induction was confirmed by western blot (fig. S1A), no growth differences were observed in either solid or liquid media (Fig. 1D and fig. S1B), suggesting that *C. tropicalis* PheRS is not the target of KLMT-1.

We next considered whether KLMT-1 may have evolved specificity for other aminoacyl-tRNA synthetases or tRNA molecules. To evaluate the global impact of KLMT-1 on tRNA biology, we performed modification-induced misincorporation tRNA sequencing (mim-tRNAseq) (*24*). This method simultaneously quantifies three key aspects of tRNA function: abundance, aminoacylation status (i.e., fraction of charged tRNA molecules), and the profile of post-transcriptional modifications, which influence tRNA stability, folding, and decoding fidelity (fig. S1C) (*25*). Consistent with our observations in *C. tropicalis* (*18*), KLMT-1 overexpression caused lethality specifically in early *C. elegans* embryos, while late embryos, larvae, and adults remained unaffected, suggesting a conserved mechanism of toxicity (Fig. 1E and fig. S1D). We therefore applied mim-tRNAseq to *C. elegans* embryos overexpressing KLMT-1 under a heat shock inducible promoter, alongside heat-shocked wild-type controls, each in biological quadruplicates (fig. S1E). However, we found no significant differences in the abundance, charging status, or modification profiles of any tRNA species—including all isoacceptors or isodecoders—between KLMT-1 expressing and control embryos (Fig. 1F,G and fig. S1F).

Altogether, these results strongly suggest that although KLMT-1 evolved from an essential PheRS subunit, it does not impair tRNA^Phe^ aminoacylation, or broadly alter the function of any tRNA species. Further, the selective sensitivity of early *C. tropicalis* and *C. elegans* embryos argues against a general disruption of mRNA translation, which would similarly impact all developmental stages. Instead, our experiments point to the existence of a distinct, conserved target—one whose function may be largely restricted to early embryogenesis.

### KLMT-1 localizes to centrosomes

Subcellular localization can offer clues about protein function. Initial attempts to visualize KLMT-1 by tagging either the intrinsically disordered N- or C-terminus with a fluorescent reporter resulted in the formation of large aggregates, precluding reliable interpretation of its native behavior *in vivo* (Fig. 1C and fig. S2A) (*18*). To circumvent this problem, we used AlphaFold3 to identify internal loops predicted to tolerate insertion of GFP without disrupting folding (fig. S2B). Guided by these predictions, we generated an endogenous *klmt-1* allele carrying an internal GFP insertion at residue Gln^234^ (hereafter KLMT-1::GFP). As expected, *klmt-1::GFP* hermaphrodites expressed KLMT-1 in the germline and the toxin was maternally loaded into unfertilized eggs (fig. S2C). Importantly, KLMT-1::GFP showed no signs of aggregation in either tissue, suggesting that the internal tag preserves its normal subcellular localization (fig. S2A). Although the endogenous *klmt-1::GFP* allele was not active in genetic crosses (fig. S2D), its lack of toxicity proved advantageous, as it allowed us to examine KLMT-1 localization in embryos lacking the antidote KSS-1, which would otherwise be dead. To this end, we generated a *kss-1* null allele in the *klmt-1::GFP* background and performed confocal microscopy of KLMT-1::GFP in both *kss-1(+)* and *kss-1*(*-*) embryos. In wild-type embryos, KLMT-1 was already efficiently degraded by the SCF^KSS-1^ complex at the late gastrulate stage, consistent with our previous observations (fig. S2E) (*18*). In striking contrast, in *kss-1*(*-*) embryos, KLMT-1::GFP concentrated into one or two discrete perinuclear foci per cell (fig. S2E). The number and position of these foci bore a striking resemblance to a hallmark feature of dividing cells: centrosomes.

Centrosomes are membrane-less highly dynamic organelles, which function as the main microtubule-organizing centers in animal cells (*26*, *27*). Beyond their central role in microtubule nucleation, centrosomes also contribute to the establishment of cell shape, polarity, and mitotic spindle assembly during cell division. At the core of each centrosome lies a pair of centrioles—barrel-shaped structures with a characteristic 9-fold microtubule symmetry. These centrioles are embedded within a much larger porous matrix known as the pericentriolar material (PCM), a protein-rich scaffold that mediates microtubule nucleation, anchoring, and organization. To determine whether KLMT-1 localizes to centrosomes, we examined its spatial relationship with established centrosomal components. To do so, we generated *C. tropicalis* reporter lines in which either *sas-4* or *spd-5* was endogenously tagged at the N-terminus with mScarlet (Fig. 2A). SAS-4 is a conserved core component of the centriole that is essential for centrosome duplication (*28*, *29*), while SPD-5 is a scaffold protein that drives PCM assembly through polymerization, serving as an organizer of PCM structure and function (*30*, *31*).

**Fig. 2.**
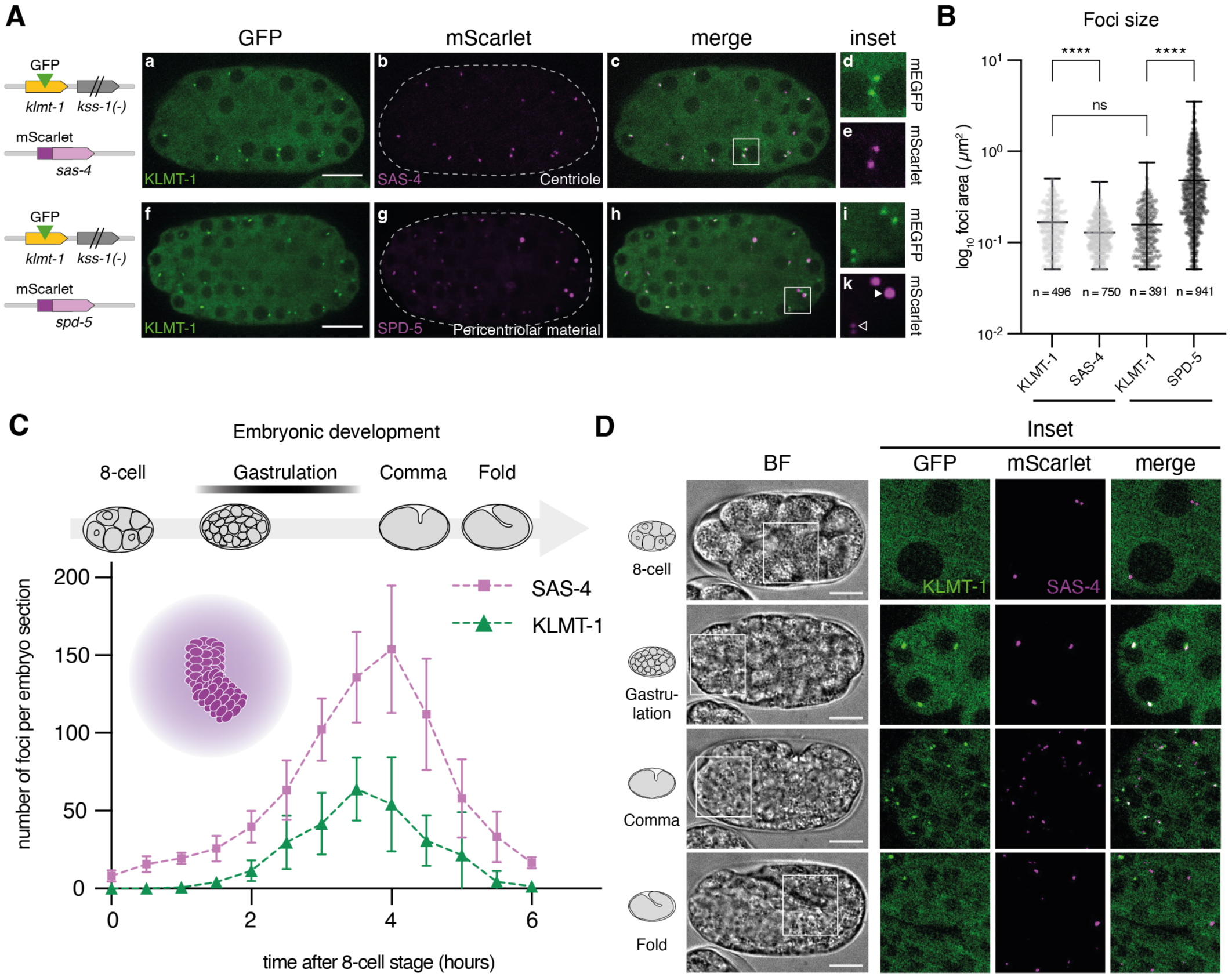
KLMT-1 localizes to centrosomes. **(A)** Co-localization of KLMT-1::GFP with known centrosomal proteins: mScarlet::SAS-4, a core centriole component (top) and mScarlet::SPD-5, a scaffold protein forming the pericentriolar material (PCM) (bottom). Scale bar is 10 μm. Insets 2.5X magnification. Arrowhead illustrates size variation of SPD-5 foci during the cell cycle. Large during mitosis (filled arrowhead) and smaller during interphase (empty arrowhead). **(B)** Quantification of foci size in *C. tropicalis* embryos expressing KLMT-1::GFP with mScarlet::SAS-4 (light grey) and mScarlet::SPD-5 (dark grey), respectively. Kruskal-Wallis test with Dunn’s multiple comparisons test. Error bars represent the mean with range. Dots represent individual foci. Data from at least 12 embryos per genetic background. **(C)** Quantification of KLMT-1::GFP and mScarlet::SAS-4 foci number per embryo during embryonic development. Mean +/- SD. Stages shown in D are indicated above. 15 embryos from 2 individual experiments were analyzed. **(D)** Representative time-lapse microscopy images of KLMT-1::GFP and mScarlet::SAS-4 foci as quantified in (C). Scale bar is 10 μm. Insets 2.0X magnification.

We next examined the SAS-4 and SPD-5 reporters in the *klmt-1::GFP; kss-1*(*-*) background using fluorescence microscopy (Fig. 2A; fig. S3A). KLMT-1 foci strongly colocalized with both SAS-4 (Manders coefficient M_KLMT-1/SAS-4_ = 0.939) and SPD-5 (M_KLMT-1/SPD-5_ = 0.945), suggesting that KLMT-1 associates with centrosomes (Fig. 2A; fig. S3B) but not all centrosomes in any given embryo co-occurred with KLMT-1 foci (mean M_SAS-4/KLMT-1_ = 0.611 and M_SPD-5/KLMT-1_ = 0.569; fig. S3B). KLMT-1 foci were on average only 25% larger than centriolar SAS-4 (166 +/- 9.3 vs. 129 +/- 5.7 nm^2^; Fig. 2B), while SPD-5 foci were substantially larger and dramatically varied in size (482 +/- 477 nm^2^, range 51-3532 nm^2^; Fig. 2B), consistent with the size fluctuations of the PCM throughout the cell cycle (*32*). By the onset of morphogenesis, most embryonic cells in *C. elegans* have exited the cell cycle and differentiated, with ∼88% of them having lost their centrioles (*33*). We observed a similar pattern in C. *tropicalis*, where coinciding with the loss of SAS-4, the vast majority of KLMT-1 foci disappeared (Fig. 2C,D). These observations indicate that KLMT-1 localizes to centrosomes, specifically in close proximity to the centrioles.

### Centrosomal recruitment of KLMT-1 is triggered in the zygote

In *C. elegans,* sperm donate centrioles at fertilization, initiating centrosome formation, which is essential for the first and subsequent cell divisions. Notably, in both *kss-1(+)* and *kss-1*(*–*) backgrounds, KLMT-1 was initially diffusely localized in the maternal gonad, the cytoplasm of unfertilized eggs and early embryos and did not co-localize with SAS-4–marked centrosomes (Fig. 2C,D; fig. S2A,C). In *kss-1(+)* embryos, KLMT-1 remained cytoplasmic throughout early development and was subsequently degraded. In striking contrast, *kss-1*(*-*) embryos exhibited a rapid relocalization of KLMT-1 to SAS-4–marked centrosomes at 100–150 min post-fertilization, immediately following the stage at which KSS-1 normally mediates its degradation (Fig. 2C,D). In contrast to KLMT-1, FARS-3 localized exclusively in the cytoplasm of cells throughout embryonic development (fig. S3D). Together, these findings suggest that KLMT-1 localization to centrosomes is not by-product of toxicity nor an ancestral feature but rather a tightly regulated process and critical for its selfish role. For a TA element to selectively kill non-carrier embryos, the toxin must be maternally deposited but not yet active prior to zygotic genome activation, allowing embryos that inherit the TA time to express the antidote and neutralize it (*16*). KLMT-1 satisfies this requirement by remaining cytoplasmic at first, allowing KSS-1–mediated degradation in TA-carrying embryos before it can associate with centrosomes, whereas in non-carrier embryos it escapes degradation and relocalizes to centrosomes. Further, centrosome targeting provides a mechanistic explanation for KLMT-1’s stage-specific lethality: it selectively kills early embryos, when centrosomes are abundant and essential, but has minimal effects on late embryos, larvae, or adults, in which centrosomes are largely absent or dispensable (*33*, *34*).

### KLMT-1 targets Aurora Kinase A

Our findings raised the intriguing possibility that KLMT-1 targets one or more proteins associated with the centrosome. A classical approach to identify such targets is through suppressor screens. Yet the narrow developmental window during which KLMT-1 is active—exclusively in early embryos—posed a significant challenge to this strategy. In pilot experiments, EMS mutagenesis followed by heat shock–induced KLMT-1 expression resulted in candidate suppressor lines. However, all proved to be false positives, likely due to the presence of a small number of late-stage resistant embryos in the population despite our best efforts. Moreover, many centrosomal genes are essential, making true suppressor alleles both rare and difficult to isolate. To circumvent these limitations, we turned to a different approach: an *in silico* pull-down screen, predicting pairwise interactions between KLMT-1 and all 106 centrosome-associated proteins known in *C. elegans* (*35*). Strikingly, while AlphaFold2 failed to detect any strong candidates, AlphaFold3 pinpointed a single, high-confidence KLMT-1 interactor: AIR-1 (Fig. 3A,B; fig. S4A,B) (*36*).

**Fig. 3.**
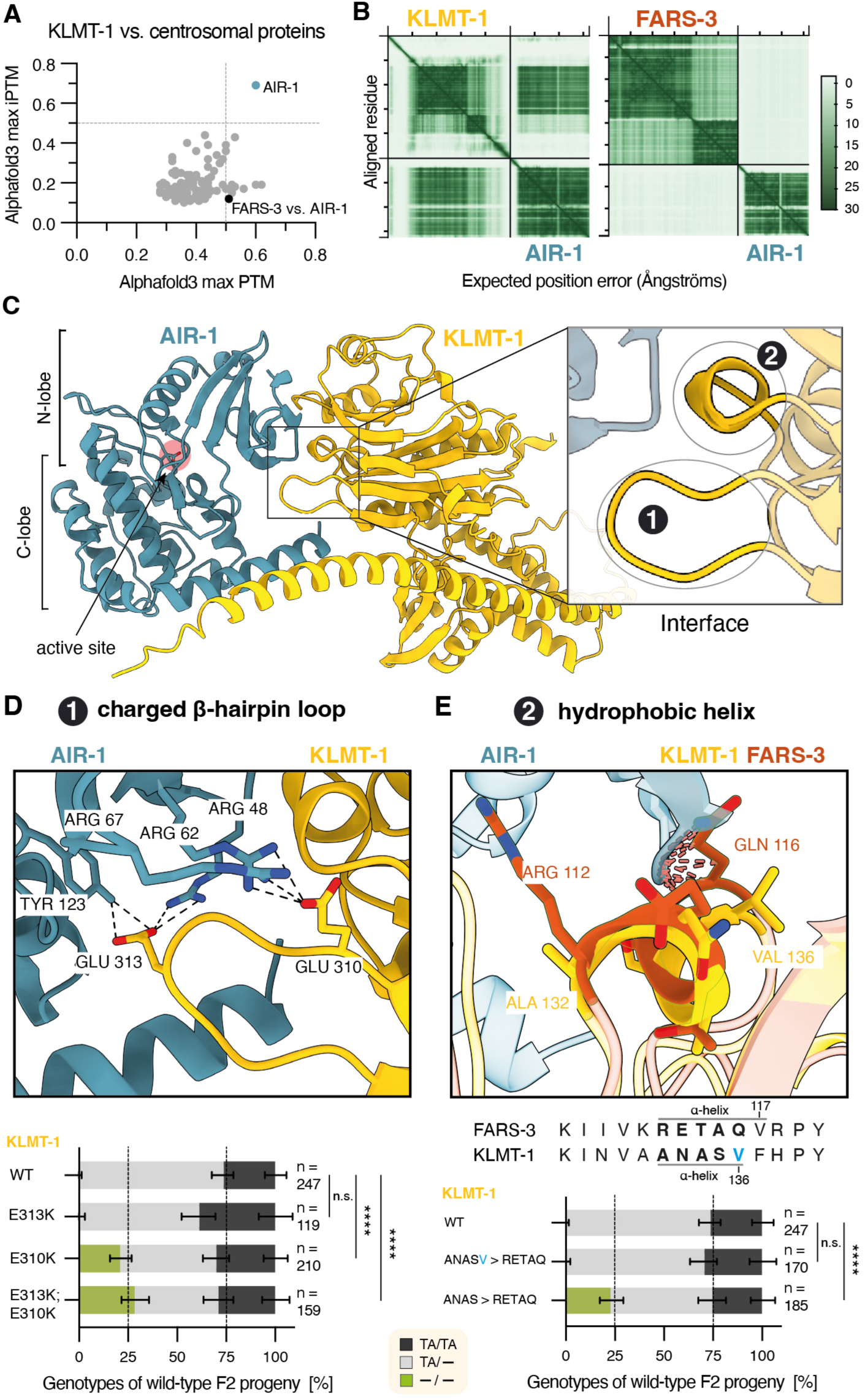
Validation of predicted bipartite interface between KLMT-1 and AIR-1. **(A)** AlphaFold3 interaction screen between KLMT-1 and all *C. elegans* proteins localized to centrosomes (GO term annotation). A prediction between *C. tropicalis* FARS-3 and AIR-1 was also included (black dot). **(B)** Predicted aligned error (PAE) plots for KLMT-1–AIR-1 (left) and FARS-3–AIR-1 (right) models. **(C)** AlphaFold3 structural model of the KLMT-1–*C. tropicalis* AIR-1 complex. Zoom-in of the KLMT-1 bipartite interface consisting of (1) a positively charged β-hairpin loop and (2) a hydrophobic α-helix contacting the N-terminal lobe of AIR-1. The active site is located in the C-lobe and is distant from the KLMT-1 interaction site. **(D)** Close-up of the predicted electrostatic interface highlighting salt bridges (top). Genetic crosses testing the effect of mutations in the KLMT-1 β-hairpin loop on toxicity (bottom). Residues Glu^310^ and Glu^313^ were mutated individually or together to lysine (Glu^310^Lys, Glu^313^Lys, and Glu^310^Lys; Glu^313^Lys) to reverse charge (Fisher’s exact test with Bonferroni correction). **(E)** Close-up of the KLMT-1 hydrophobic helix predicted to contact AIR-1, compared with the equivalent FARS-3 helix. Dashed lines indicate steric clashes predicted between AIR-1 and bulkier ancestral FARS-3 residues, including Arg^112^ (corresponding to Ala^132^ in KLMT-1) and Gln^116^ (corresponding to Val^136^ in KLMT-1) (top). Sequence alignment showing the KLMT-1 hydrophobic helix and equivalent FARS-3 residues (middle). Genetic crosses testing the effect of reverting KLMT-1 helix residues to FARS-3-like identities (bottom). Only insertion of the sterically more constrained ANAS helix abolishes toxicity (*n*_(-/-)_ = 42/185, *P*_adj_ < 0.0001, Fisher’s exact test with Bonferroni correction).

AIR-1 is the ortholog of Aurora kinase A, a highly conserved serine/threonine kinase from yeast to humans (*37*). Aurora kinase A localizes to centrosomes during interphase and is required for mitotic entry, centrosome maturation and separation, and chromosome alignment (*38–42*). Consistent with the specificity of this interaction, *C. tropicalis* FARS-3 was not predicted to bind AIR-1 (Fig. 3A,B). AlphaFold3 indicated that KLMT-1 engages AIR-1 via a bipartite interaction surface consisting of a negatively charged surface loop that inserts into a positively charged pocket on AIR-1’s N-terminal lobe and a hydrophobic α-helix in close spatial proximity (Fig. 3C; fig. S4C). The loop, which connects two antiparallel β-sheets, contains two glutamic acid residues, Glu^310^ and Glu^313^, predicted to form salt bridges with AIR-1 residues Arg^62^ and Arg^48^, respectively (Fig. 3D). To validate this interaction *in vivo*, we first introduced two charge-reversal mutations, Glu^310^Lys and Glu^313^Lys, in the endogenous *klmt-1* locus using CRISPR/Cas. These two mutations did not affect KLMT-1 expression levels (fig. S4D); however, genetic crosses to the susceptible strain revealed that KLMT-1 toxicity was fully abrogated, as evidenced by restored Mendelian segregation in the F_2_ generation (Fig. 3D). In addition, we generated single mutations and found that the Glu^310^Lys charge reversal was sufficient to fully abolish KLMT-1 toxicity (Fig. 3D). We next asked whether the adjacent hydrophobic α-helix also contributes to this interaction (Fig. 3C and E). In FARS-3, the corresponding helix is predicted to be one residue longer and to contain bulkier side chains, such as Gln^116^, which could introduce steric clashes with AIR-1 (Fig. 3E). To assess the functional relevance of these changes, we replaced the KLMT-1 helix with its FARS-3 counterpart using two strategies. In the first, we performed a length-preserving substitution (KLMT-1 ‘ANASV’ to FARS-3 ‘RETAQ’) assuming one-to-one equivalence of all residues. In the second, we replaced the helix while extending it by one residue (KLMT-1 ‘ANAS’ to FARS-3 ‘RETAQ’). This strategy preserved KLMT-1 last helical residue, Val^136^, and more-closely matched the predicted ancestral architecture (Fig. 3E). While the length-preserving substitution had no detectable effect on KLMT-1 toxicity, replacing it by the longer, FARS-3–like helix completely abolished toxicity (Fig. 3E, fig. S4D). These results suggest that the shorter, less bulky KLMT-1 helix is important for productive binding to AIR-1, likely by minimizing steric constraints at the interface compared to FARS-3.

### A mutation in Aurora Kinase A suppresses KLMT-1 toxicity

The evidence presented thus far grants KLMT-1 with both the opportunity and the means—spatial proximity and structural compatibility—for interacting with AIR-1. However, it does not formally show that AIR-1 is its direct target. To show this, we set out to suppress KLMT-1 toxicity by mutating the predicted AIR-1 interface. This approach, however, presented significant challenges. Unlike *klmt-1*, which is dispensable for embryonic development, *air-1* is both essential and highly conserved in eukaryotes (Fig. 4A) (*39*). For instance, Arg^62^, the key negatively charged residue predicted to form salt bridges with KLMT-1, is conserved from yeast to humans (Fig. 4A). Moreover, the predicted KLMT-1 binding site on AIR-1 overlaps with the interface used by known AIR-1 allosteric regulators, complicating efforts to selectively disrupt the interaction without hindering AIR-1 activity and killing the worms in the process (Fig. 4B)

**Fig. 4.**
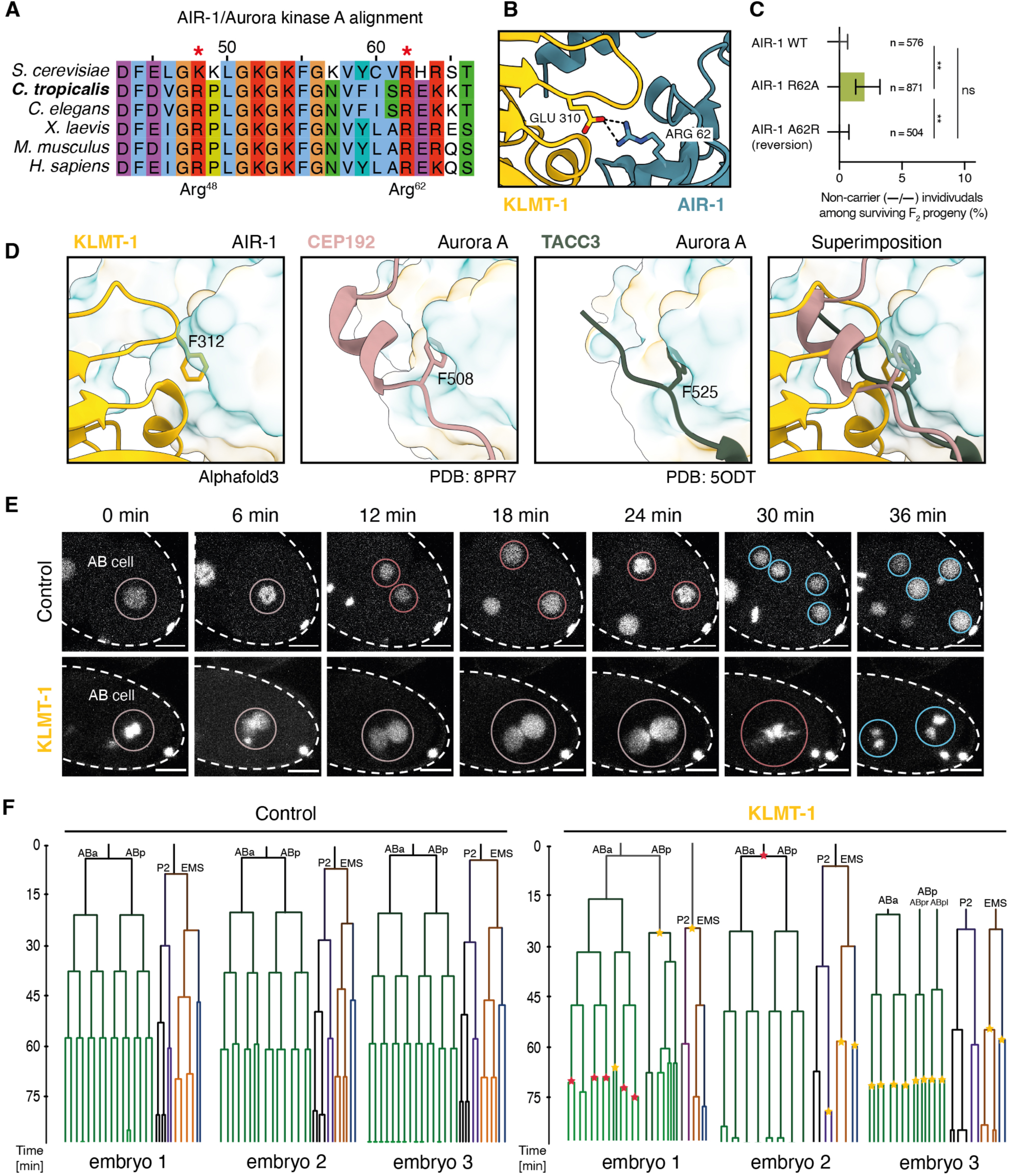
An AIR-1 mutation suppresses KLMT-1 toxicity, which otherwise causes cell-cycle and chromosome segregation defects. **(A)** Multiple sequence alignment of AIR-1 (Aurora kinase A) orthologs across species showing strong conservation of two positively charged residues, Arg^48^ and Arg^62^ (red asterisks). **(B)** Predicted KLMT-1–AIR-1 interaction interface highlighting a putative salt bridge between KLMT-1 Glu^310^ and AIR-1 Arg^62^ (left). **(C)** Genetic crosses between the *klmt-1/kss-1* TA carrier and a susceptible strain in three genetic backgrounds: *air-1*(wt), *air-1*(Arg^62^Ala), and the reversion mutant *air-1*(Ala^62^Arg). Wild-type (-/-) progeny are only observed in the R62A background (*n* = 18/871, *P*_adj_ = 0.0009, Fisher’s exact test with Bonferroni correction). **(D)** Comparison of the predicted KLMT-1–AIR-1 binding interface with solved structures of human Aurora-A bound to known allosteric regulators CEP192 (PDB 8PR7) and TACC3 (PDB 5ODT). Convergent phenylalanine residues mediating hydrophobic docking into the kinase N-terminal lobe are highlighted. **(E)** Representative time-lapse images showing early division defects in the AB cell lineage upon KLMT-1 overexpression compared with control embryos. Nuclei are visualized using histones HIS-58 and HIS-24 fused to mCherry fluorescent reporter. **(F)** Representative lineage trees from three KLMT-1 overexpression embryos and three wild-type embryos (left). Trees were temporally aligned to the ABa division, with the time scale shown on the left. Early embryonic cell names are labeled. Only the first abnormal division event in each branch is marked, delayed and early divisions are indicated by yellow and red stars, respectively.

Despite these constraints, we proceeded with a targeted CRISPR-editing strategy. As with KLMT-1, we introduced two charge-reversal mutations in AIR-1, Arg^48^Glu and Arg^62^Glu, designed to disrupt predicted salt bridges with KLMT-1’s negatively charged glutamic acid residues (Fig. 4B,3C). Unfortunately, both the double mutant, as well as individual Arg^48^Glu and Arg^62^Glu single mutants proved to be homozygous lethal, preventing us from testing whether they could suppress KLMT-1 toxicity (fig. S5A). Reasoning that a charge-neutral substitution might be better tolerated than a full reversal, we next generated an AIR-1 Arg^48^Ala; Arg^62^Ala double mutant. Yet again, the result was homozygous lethality, reinforcing just how crucial these residues are for AIR-1 function. Next, we generated Arg^48^Ala and Arg^62^Ala single charge-neutralization mutants. The first one was once again lethal but to our fortune, the Arg^62^Ala single mutant (fig. S5A,B) could be maintained in the homozygous state. Although AIR-1 Arg^62^Ala worms exhibited high lethality and multiple fitness defects, approximately 25% of individuals developed into fertile adults (fig. S5C), providing a valuable foothold for probing AIR-1’s contribution to KLMT-1 toxicity.

To test whether mutating the predicted AIR-1 interface could suppress KLMT-1 toxicity, we crossed *klmt-1/kss-1* TA carrier and non-carrier strains in both *wt* and *air-1* (Arg^62^Ala) backgrounds. In the wild-type background, once again, and consistent with KLMT-1’s potent toxicity, we did not recover a single homozygous non-carrier survivor among more than 491 individually genotyped F_2_ progeny, indicating that spontaneous escapers are exceedingly rare, if they occur at all (Fig. 4C). In contrast, introduction of the *air-1* (Arg^62^Ala) mutation led to reproducible recovery of homozygous non-carrier progeny: 18 such individuals were identified among 833 F_2_ animals (2.16% of total F_2_, compared to a Mendelian expectation of 25%, Fig. 4C). This increase in non-carrier recovery relative to wild type was highly significant (Fisher’s exact test, *P*adj = 0.0009). When normalized to the expected Mendelian frequency, this corresponds to 8.6% rescue of the homozygous non-carrier class (95% CI: 5.5–13.6%), indicating partial but robust suppression of KLMT-1 toxicity. This incomplete suppression is consistent with AlphaFold3 predictions suggesting that the Arg^62^Ala substitution alone is not sufficient to fully abolish binding between AIR-1 and KLMT-1 (fig. S5D). To confirm that the suppression effect was specifically caused by the *air-1* (Arg^62^Ala) mutation and not by a secondary mutation that indirectly reduced KLMT-1 toxicity, we used CRISPR/Cas to revert *air-1* Ala^62^ back to the wild-type Arg residue. Genetic crosses in the repaired *air-1* (Ala^62^Arg) background were indistinguishable from wild type: no homozygous non-carrier survivors were detected among 572 genotyped F_2_ individuals (Fig. 4C). Together, these results indicate that mutation of AIR-1 Arg^62^specifically suppresses KLMT-1 toxicity and strongly support a model in which KLMT-1 exerts its toxic effect through direct interaction with AIR-1, mediated by KLMT-1 Glu^310^ and AIR-1 Arg^62^.

### KLMT-1 phenocopies loss of AIR-1

The activity of AIR-1 is tightly regulated by a dynamic network of allosteric modulators that control its kinase activity, as well as localization to centrosomes and mitotic spindles (*43*). Key regulators include SPD-2 (Cep192), TPXL-1 (Tpx2), TAC-1 (TACC3) and SPAT-1 (BORA) (*44–48*). Structural comparisons between human Aurora Kinase A in complex with its regulators and the predicted AIR-1–KLMT-1 binding interface revealed a substantial overlap and convergence of their interaction surfaces (Fig. 4D). For example, human Aurora Kinase A residue Arg^151^, corresponding to AIR-1 Arg^62^, forms stabilizing salt bridges with CEP192 (*49*). Further, the KLMT-1 β-hairpin loop contains a phenylalanine (Phe^312^) predicted to insert into a hydrophobic pocket of AIR-1, mirroring the binding mode of canonical Aurora interactors (Fig. 4D). Together, these observations suggest that KLMT-1 may inhibit AIR-1 activity by competing for binding with its known allosteric modulators. In *C. elegans*, a*ir-1* RNAi treatment results in arrested embryos that are severely aneuploid, including cells with large polyploid nuclei. Early embryos display incomplete pronuclear fusion, anaphase chromatin bridges, and widespread chromosome segregation defects (*39*). Based on these observations, we hypothesized that KLMT-1 overexpression would phenocopy AIR-1 depletion and lead to various downstream cell-cycle and chromosome segregation defects. To test this prediction, we turned to *C. elegans* due to its fully resolved cell lineage and advanced single-cell tracking tools (*50*, *51*).

We induced a single-copy KLMT-1 transgene in the germline of mothers using a heat-shock promoter, collected embryos, and performed live time-lapse imaging of fluorescently labeled histones to reconstruct cell lineages (n=21) (*52*) (see Materials and Methods). KLMT-1 induction caused a broad spectrum of developmental defects that closely resembled those reported following *air-1* RNAi (*39*). These defects included chromosome segregation defects, abnormal nuclear content, cell-cycle timing defects, and cell displacement through embryogenesis (Table 1, Fig. 4E-F, Movies S1-6, and fig. S6A). In contrast, wild-type embryos expressing only fluorescently labeled histones developed normally under both control (n=8/8) and heat-shock conditions (n=10/11).

**Table 1.** Summary of KLMT-1–associated embryonic phenotypes. Embryos were scored for the presence or absence of phenotypes. Values are shown as the number of embryos displaying each phenotype over the total number analyzed (n/N with percentages). Phenotypes were not mutually exclusive. WT, wild type; HS, heat shock.

| Phenotype category | WT | WT + HS | KLMT-1 | KLMT-1 + HS |
| --- | --- | --- | --- | --- |
| Chromosome segregation defects | 0/8 (0%) | 1/11 (9.1%) | 0/7 (0%) | 9/21 (42.9%) |
| Abnormal nuclear content | 0/8 (0%) | 0/11 (0%) | 0/7 (0%) | 10/21 (47.6%) |
| Cell cycle timing defects | 0/8 (0%) | 1/11 (9.1%) | 0/7 (0%) | 11/21 (52.4%) |
| Abnormal cell displacement | 0/8 (0%) | 0/11 (0%) | 1/7 (14.3%) | 11/21 (52.4%) |

Prior to the first mitotic division, KLMT-1–overexpressing embryos frequently displayed supernumerary or irregular nuclei, consistent with failures in pronuclear fusion or incomplete meiotic chromosome segregation (n =4/10). All embryos imaged from the zygotic stage onward exhibited an abnormal first division (6/6). Notably, the early defects observed here differ from the phenotype in *C. tropicalis* crosses, where endogenous KLMT-1 neither perturbs the first embryonic divisions nor is yet localized to centrosomes. Given that AIR-1 is already present at centrosomes from the first cell division, this discrepancy is unlikely to reflect differences in target availability (*39*). Instead, elevated KLMT-1 levels resulting from transgenic overexpression likely drive premature engagement with AIR-1. Alternatively, the premature phenotype may reflect species-specific differences in sensitivity. Nonetheless, our results indicate that KLMT-1 overexpression disrupts mitosis and recapitulates downstream consequences of AIR-1 depletion, consistent with impaired AIR-1 activity in vivo.

### Evolution of the KLMT-1 binding interface

Although the Glu^310^ residue in the KLMT-1 β-hairpin is essential for KLMT-1 toxicity, an equivalent glutamic acid, Glu^292^, is also present in *C. tropicalis* FARS-3. Despite this, AlphaFold3 did not predict an interaction between FARS-3 and AIR-1 (Fig. 3B). Structural analysis of published models and AlphaFold3 predictions suggested that this discrepancy arises from differences in the length of the loop of the β-Hairpin motif: *C. tropicalis* FARS-3 contains a shorter loop than KLMT-1 (Fig. 5A). As a result, the three glutamic acids in the FARS-3 loop are neither correctly positioned nor oriented for interacting with AIR-1. A multiple sequence alignment of KLMT-1 and FARS-3 orthologs suggested that this structural difference likely arose from a three–amino acid insertion downstream of Glu^310^—a change that extended the KLMT-1 loop and reoriented its side chains for optimal AIR-1 engagement (Fig. 5B). To directly test this evolutionary model, we reverted the KLMT-1 loop to a FARS-3–like configuration while preserving the key glutamic acid residue, Glu^310^ (Fig. 5A,B). Specifically, we replaced Pro^315^ with the ancestral lysine and deleted residues 316–318 at the endogenous *klmt-1* locus. This reversion (Pro^315^Lys; ΔRAL^316–318^) did not affect KLMT-1 expression levels but completely abolished its toxicity in genetic crosses (Fig. 5B, fig. S4C). Strinkly, incorporating either the FARS-3-like loop reversal or the Glu^310^Lys charge reversal mutation in the KLMT-1::GFP construct abolished KLMT-1 centrosomal localization, strongly suggesting that interaction of KLMT-1 with AIR-1 is critical for either recruitment or maintenance of the toxin in these organelles (Fig. 5C,D; fig. S7A).

**Fig. 5.**
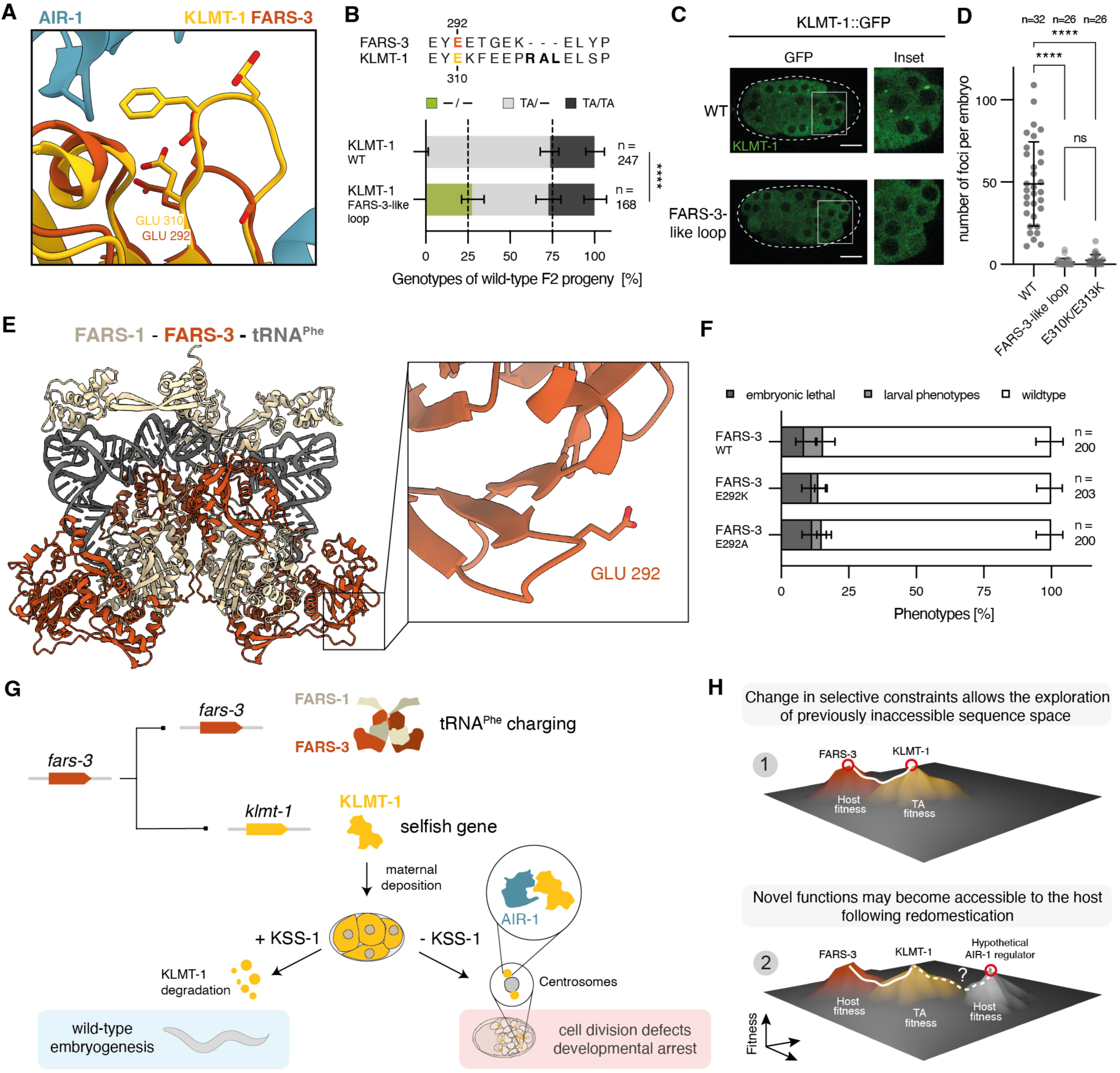
The KLMT-1 toxic interface evolved from a largely neutrally-evolving linker of FARS-3. **(A)** Predicted KLMT-1–AIR-1 interaction interface. The FARS-3 structure is superimposed onto KLMT-1 to compare their homologous β-hairpin loop regions. Glu^310^ in KLMT-1 is essential for toxicity, and the equivalent residue (Glu^292^) is conserved in FARS-3. **(B)** Sequence alignment of FARS-3 and KLMT-1 showing a three–amino acid insertion that extends the KLMT-1 β-hairpin loop (top). Genetic crosses testing the effect of reverting this loop to an ancestral FARS-3–like state on KLMT-1 toxicity (bottom). KLMT-1 with a FARS-3-like loop is fully non-toxic (*n*_(-/-)_ = 46/168, *P* < 0.0001, Fisher’s exact test). **(C)** Single-copy KLMT-1::GFP reporter imaging showing how toxicity-abolishing mutations affect KLMT-1 centrosomal localization. Scale bar = 10µm. **(D)**Wild-type KLMT-1 localizes to the cytoplasm and centrosomes, whereas ancestral loop-reversion (middle) or charge-reversal (right) mutants show cytoplasmic localization only (*P* < 0.0001, one-way ANOVA with Tukey’s multiple comparisons test). Mean +/- SD. Dots represent individual embryos from 2 experiments. **(E)** AlphaFold3 model of the *C. tropicalis* PheRS complex containing two FARS-1 (beige) and two FARS-3 (orange) subunits bound to two tRNA^Phe^ molecules (grey). Inset highlights FARS-3 Glu^292^ residue, which is solvent-exposed and does not contact other complex components. **(F)** Phenotypes of wild-type worms compared with homozygous *fars-3*(Glu^292^Lys) and *fars-3*(Glu^292^Ala) mutants. No significant differences were observed (Fisher’s exact test). **(G)** Evolutionary origin and toxicity mechanism of KLMT-1. KLMT-1 originated from the tRNA synthetase subunit FARS-3 through gene duplication and is maternally deposited into embryos. In the presence of the antidote KSS-1, KLMT-1 is efficiently degraded, allowing normal embryonic development. In the absence of KSS-1, KLMT-1 relocalizes to centrosomes, where it interacts with AIR-1, leading to mitotic defects and ultimately developmental arrest. **(H)** Model illustrating how toxin–antidote (TA) elements may facilitate evolutionary innovation. The sequence space accessible to the essential protein FARS-3 (orange fitness peak) is constrained by strong purifying selection imposed by organismal fitness requirements. Gene duplication and altered selective pressures allow the duplicated copy to explore previously inaccessible regions of sequence space (yellow fitness peak). Subsequent domestication and functional reintegration into host biology may enable the emergence of novel adaptive functions (white fitness peak).

In agreement with its universally essential role, FARS-3 is a highly conserved protein. Within *Caenorhabditis* nematodes, the key β-hairpin loop displays little sequence variation and retains Glu^292^, corresponding to KLMT-1 Glu^310^, along with two additional glutamic acids that together confer a net negative charge (fig. S7B). However, across more distantly related taxa, this loop has diverged substantially, and neither the presence of Glu^292^ nor its overall negative charge is conserved (fig. S7B). This degree of variability is consistent with the loop’s solvent-exposed position and with structural data indicating that it does not contact either FARS-1 or tRNA^Phe^ (Fig. 5E) Supporting this view, charge-neutralizing (Glu^292^Ala) or charge-reversing (Glu^292^Lys) mutations introduced at the endogenous *fars-3* locus produced homozygous viable lines indistinguishable from wild-type, indicating that Glu^292^ is dispensable for tRNA^Phe^ charging (Fig. 5F). Together, these findings suggest that following *fars-3* duplication, KLMT-1 acquired a novel bipartite AIR-1 binding interface through two key evolutionary changes: (i) a three–amino acid insertion that extended the β-hairpin loop and repositioned a key ancestral glutamic acid, and (ii) additional substitutions that rendered a neighboring helix less bulky, thereby avoiding steric clashes with AIR-1 (Fig. 3D,E). Lastly, our results strongly suggest that the evolution of this radically novel function was contingent on substitutions that accumulated neutrally in FARS-3 during nematode evolution.

## Discussion

Deep learning–based generative models have revolutionized structural biology. While numerous studies have demonstrated AlphaFold’s ability to accurately model highly conserved complexes (*53–55*), its ability to predict interactions in the absence of strong co-evolutionary signals remains unclear. Here, we report and validate what is, to our knowledge, the first example of AlphaFold3 identifying a high-confidence interaction restricted to a single species. This observation may reflect the shift from the AlphaFold2 *evoformer* to the *pairformer* module, which reduced reliance on multiple sequence alignments. More broadly, our findings highlight the potential of such deep-learning models to explore highly specific and phylogenetically restricted protein complexes.

Aurora kinase regulation presents a paradox: multiple, distinct proteins have independently evolved to bind the same regulatory interface (*46*, *47*, *49*, *56*). Such convergence suggests the existence of mechanisms that facilitate exploration of this shared interaction space. Inspired by this evolutionary pattern, and supported by our results, we propose that selfish TA systems may act as evolutionary incubators for the emergence of new protein functions (Fig. 5G). In nematodes, plants, fungi, and vertebrates, TA systems originate from host genes through gene duplication (*16*). The transition of a duplicated host gene into a selfish toxin entails a fundamental shift in selective constraints: rather than being optimized for organismal fitness, toxins are selected to exploit vulnerabilities in host biology to ensure their own propagation. This shift both relaxes constraints and imposes new selective pressures, allowing the exploration of sequence and functional space that would be inaccessible in the ancestral context (Fig. 5G).

The emergence of an AIR-1 inhibitor from an essential tRNA synthetase subunit illustrates how such a shift can give rise to new functions, which may subsequently be refined and co-opted back into host biology through domestication, for example following duplication of the toxin (Fig. 5G). We therefore propose that some host proteins may represent the stabilized end products of such host–selfish–host evolutionary cycles (Fig. 5G) with innovation unfolding in a transient fast-paced phase that is rarely preserved, leaving its impact on evolution largely underestimated. While this framework may not necessarily explain the origin of known Aurora kinase regulators, it provides a general mechanism by which proteins can explore otherwise inaccessible regions of sequence space, thereby facilitating the emergence of new from old.

## Acknowledgments

We thank members of the Burga for their feedback and comments on the manuscript. We thank members of the Dammermann, Gerlich, Brennecke and Matos labs for reagents and advice on experiments.

## Author contributions

A.B. conceived and supervised the project. J.J.R. and A.B. designed and developed the project. J.J.R., P.T., and K.A.E. performed crosses and molecular genetics work. A.H., J.J.R., and G.D. performed structure-guided analyses and experimental validations. D.K. performed yeast experiments. J.G. and P.D. designed and generated transgenic worm lines. K.X and C.V.B. performed lineaging experiments and analyses supervised by R.J. mim-tRNAseq was performed by T.B. supervised by D.N. A.B. and J.J.R. wrote the manuscript.

## Data and material availability

All raw sequencing data and genome assemblies are available under National Center for Biotechnology Information (NCBI) project no. PRJNA1457981.

## Supplementary Materials

Figs. S1 to S7

Tables S1 and S4

Movies S1 to S6

Data S1 to S5

## Materials and Methods

### Maintenance of worm strains

Nematodes were grown on modified nematode growth medium (NGM) plates with 1% agar/0.7% agarose to prevent *C. tropicalis* burrowing. Plates were seeded with *E. coli* OP50. Experiments with *C. tropicalis* were conducted at 25 °C, whereas *C. elegans* experiments were performed at 20 °C. In case of bacterial contamination, worms were bleached using 500 µl of 1:1 mix of 1M NaOH and 5% bleach added to 750 µl of worms in M9, bleaching was stopped by washing embryos twice in 14 ml of M9. Supplemental Table S1 lists all the strains used in this study, some of which were provided by the CGC, funded by the NIH Office of Research Infrastructure Programs (P40 OD010440).

### Phenotyping and genotyping of crosses

For crosses, 4–5 L4 hermaphrodites were mated with 25–50 males (1:6 - 1:10 ratio) in 5 cm plates with modified NGM. After two days 15 L4 F_1_ progeny were transferred in bulk to a fresh plate, and then the next morning singled into individual plates, to ensure synchronized laying of F_2_ generation embryos. After 2-3 hours of active laying F_1_ hermaphrodites were collected for genotyping by PCR, and at least 10 embryos from each heterozygous hermaphrodite were transferred to individual plates the same day. F_2_ were staged and phenotyped daily until they either died or reached sexual maturity. Embryonic lethality, larval lethality, arrested development, delayed reproduction and sterility were assessed. In all crosses, we assessed the distribution of genotypes among wild-type F_2_ progeny as a measure of KLMT-1 function. For genotyping by PCR worms were digested with lysis buffer (10 mM Tris-Cl pH8, 50 mM KCl, 2.5 mM MgCl_2_, 200 µg/ml Proteinase K). PCRs were performed using in-house hot-start Taq polymerase (Molecular Biology Service, IMBA). If necessary, PCR products were additionally analyzed by Sanger sequencing. Efficiency of PCR was low for progeny arrested as embryos or at early larval stages. Detailed information on genotyping and phenotyping of crosses can be found in Supplemental Data S1.

### Phenotyping of *C. tropicalis* strains

Around 15 L4 larvae were picked one day prior to the experiment to ensure a homogeneous population of young adults. On day 0, around 10 adult worms were singled and allowed to lay for 1-2 hours. Approximately 10 embryos per adult were then transferred in bulk to fresh 5 cm modified NGM plates with OP50. Development of embryos at 25°C was tracked for 3 days, scoring embryonic and larval lethality, delay or infertility. Wild-type worms were egg-laying adults (or, in rare cases, adult males) on day 3. All phenotyping was performed at least in duplicates.

### Wild-type-only bulk crosses

In cases I) of known detailed phenotypic outcome of a cross and/or II) where high background lethality would otherwise prevent obtaining sufficient numbers of wild-type cross progeny, we employed a modified cross protocol in which singled F_1_ mothers were allowed to lay for an extended period of time (16-24h) prior to genotyping. Subsequently, the F2 plates from confirmed heterozygous mothers were incubated for three days. Only wild-type F2 progeny (gravid adults and adult males) were collected individually and genotyped as described above. Detailed information on genotyping and phenotyping of crosses can be found in Supplemental Data S1.

### Generation of transgenic lines

For CRISPR–Cas gene editing, we adapted previously described protocols (*57*). In brief, 250 ng/µL Cas9 or Cas12a proteins (IDT) were incubated with 200 ng/µL crRNA (IDT) and 333 ng µl−1 tracrRNA (IDT) at 37 °C for 10 min before adding 2.5 ng/µL co-injection marker plasmid (pCFJ90-mScarlet-I, or pCFJ90-mNG for editing of lines expressing mScarlet-tagged proteins). For homology-directed repair, donor oligos (IDT) or biotinylated and melted PCR products were added at a final concentration of 200 ng/µL or 100 ng/µL, respectively. Following injections into young hermaphrodites, mScarlet or mNG-positive F_1_ progeny were singled and their offspring screened by PCR and Sanger sequencing to detect successful editing. To insert single-copy transgenes we adapted a split selection system developed for *C. elegans* (*58*). We introduced a synthetic landing pad (SLP) with split hygromycin B resistance in Chr. IV of *C. tropicalis* EG6180. For homology-directed repair, we used plasmids carrying an insert of interest followed by the N-terminal portion of the hygromycin resistance gene under ribosomal promoter *Ctr*-*rps-20*p, at a final concentration of 50–60 ng/µL. On day 3 or 4 post-injection, we supplemented plates with 600 µL of 5 mg/mL hygromycin B. Then, 7–10 days after poisoning, survivors were singled, propagated and screened by PCR and Sanger sequencing to detect successful editing. All guide RNAs and HDR templates as well as plasmids used for SLP injections are listed in Supplementary Table S2 and S3 and are available upon request.

### Heat shock of *C. tropicalis* embryos and larvae

Heat-shock induction was performed on 5-cm plates with modified NGM. For embryonic stages, embryos were transferred to the plates after dissecting gravid adults with insulin syringe needles (29 G) in a drop of M9 and were heat shocked in a 37 °C incubator for 40 min, either after briefly drying (early embryos; on average 25 min) or after incubation for 6h (late embryos; corresponding to development until comma stage). For larval stages (L1/L4), worms were transferred to plates, briefly dried and heat shock treated as described above. During heat shock, plates were kept with their lids on top as opposed to standard upside-down plate storage, to ensure even warming. The number of embryos or larvae was manually counted after heat-shock induction, and plates were kept in a 25 °C incubator overnight. Wild-type, unhatched or affected embryos or larvae were tracked for two days with daily documentation. The percentage of affected worms was calculated using a sum of non-hatched embryos and affected larvae divided by the total number of embryos/larvae used for heat shock for each strain. All experiments were performed in triplicates. Raw numbers of heat shock phenotyping can be found in Supplemental Data S2. Induction of KLMT-1 after heat shock was validated by western blot (fig. S1D).

### Yeast genetics and Ctr-PheRS heterologous system

The temperature sensitive FRS2 S. cerevisiae strain (YFL022C) (*23*) was transformed with pCEV-G2-Km::*fars-1*::*fars-3* or with pCEV-G2-Km::*fars-1*::*fars-3* and p42Nat::pGal1-10::*klmt-1* and spread on selective YPDA plates (200 µg/mL G418; 100 µg/mL CloNat) at 30°C for 3-5 days. For spot assays, yeast cultures were prepared by streaking freshly transformed YFL022C onto YPD agar plates containing antibiotics (CloNat 100 µg/mL; G418 200ug/ml). After incubating the cells at 25°C in YPD supplemented with 2% Glucose overnight, cultures were resuspended in autoclaved H2O to a final OD600 of 1.0. Serial dilutions (10^−1^ to 10^−6^) were prepared by mixing 10 μL of culture into 90 μL sterile H_2_O. From each dilution, 5 μL was spotted in sequence onto solid agar plates with varying composition depending upon the strain and experimental design. Plates were incubated at 25°C or at 37°C for 48 h and scanned for growth comparison. For liquid culture growth assays, synthetic defined minimal media (SDMM) was made using dropout base powder (SC-Phe, Sunrise Science) and M9 minimal broth. For M9 minimal broth 200 mL 5x M9 Salts (5x M9 Salts composed of 64g Na_2_HPO_4_, 15g KH_2_PO_4_, 2,5 g NaCl, 5g NH_4_Cl per Liter) was prepared and the following filter sterilized solutions were added: 2 mL 1 M MgSO4 solution, 0.1 mL 1 M CaCl_2_ solution and 100 mL 20% glucose, and filled up to 1000 mL with sterile MonoQ. For SDMM containing 2% raffinose and 2% raffinose / 2% galactose, 100 mL of 20% raffinose or 50ml of 40% raffinose/galactose was added. For all growth assays, S. cerevisiae strains and derivatives were streaked on YPDA and grown at 25°C for 72 h. Single colonies were suspended YPD with 2% glucose for overnight culture. Overnight cultures were regrown in fresh YPD with 2% glucose media for several hours until reaching an OD600 of 0.1-0.3. Cultures were washed with sterile MonoQ and used to inoculate 200 μl of media in a microtiter plate to a final OD600 of 0.01. Growth was monitored using a Synergy H1 absorbance spectrophotometer (BioTek) by measuring the absorbance at 600 nm every five minutes for 72 h. Rescue of FRS2 strains was achieved through transformation of plasmid pCEV containing C. *tropicalis* FARS-1 and FARS-3.

### Modification-induced misincorporation tRNA sequencing (mim-tRNAseq)

For sample preparation, approximately 40 000 synchronized L1 larvae were seeded on NGM plates with *E. coli* OP50 and incubated at 20°C for 3.5 days, until predominantly young gravid adults and laid embryos were present. Gravid adults were collected in 15 mL Falcon tubes and sedimented to minimize carryover of laid embryos. Samples were bleached as described above, heat shocked on unseeded prewarmed 15 cm NGM plates (1h, 37°C), and incubated for 5h at 20°C. Embryos were collected in M9, approximately 1/5 of the sample was removed for protein extraction and western blot analysis, while the remaining sample was flash-frozen in TRIzol (Invitrogen, 15596018) and stored at -80°C until further processing. Samples were processed for tRNA sequencing as described in Behrens et al (*59*) with minor modifications. Briefly, RNA was extracted and precipitated on ice under mildly acidic conditions (using chloroform instead of BCP). Following oxidation, β-elimination and dephosphorylation, RNA was size-selected (60-100 nt) by excision and elution from a denaturing polyacrylamide gel. Following 3’ adapter ligation and subsequent polyacrylamide gel purification, the RNA was reverse transcribed using Induro RT (NEB), which is reported to have comparable performance to the discontinued TGIRT enzyme (*60*). Following template hydrolysis and gel purification, the obtained cDNA was circularized with CircLigase (Lucigen) and libraries were prepared by 5’/3’ index primer addition and amplification with KAPA HiFi DNA polymerase (Roche). The resulting libraries were concentrated, quantified, and sequenced on an Illumina NovaSeq6000 sequencing system. Following demultiplexing and adapter trimming, tRNA expression, charging and modifications were analyzed with the mim-tRNAseq computational package (v1.3.8) using the following command: mimseq --species Cele - -cluster-id 0.9 --threads 40 --control-condition N2 --max-mismatches 0.1 --remap-mismatches 0.075 -n Celeg_KLMT --remap --deconv-cov-ratio 0.4. Abundance, charge and modification data from tRNA sequencing used for Fig. 1/fig. S1 can be found in Supplemental Data S3.

### Imaging of developing embryos and cell lineaging

*C. elegans* embryos were collected in M9 medium with 20 μm beads and mounted under a coverslip as described previously (*51*). Imaging was performed using a Zeiss LSM 880 confocal microscope with a 63X oil immersion objective. Nuclei were visualized using an integrated mCherry tagged histone marker expressed in the RW10226 strain. Time-lapse z-stack recordings were acquired at 90 seconds intervals for up to 2 hours, and embryos were analyzed until approximately the fourth division of the AB cell. Cell lineages were traced using a semi-automated pipeline developed in-house. A U-Net was trained on a manually annotated ground-truth dataset of nuclear z-stacks to generate 3D instance segmentations of nuclei across timepoints. Due to the limited size of the training data and inherent noise in live imaging, segmentation quality required systematic manual correction. To facilitate this, a dedicated GUI application was developed for visualizing and editing lineage data, allowing users to scroll through image and segmentation volumes, and to add, remove, or modify cell identities, connections, and division events. All lineages were manually reviewed and corrected with this tool prior to downstream analysis. The tool is available at: https://bitbucket.org/pgmsembryogenesis/lineager. Cell-cycle timing defects were defined as deviations greater than 3σ from the wild-type mean division timing, where the wild-type mean was estimated by MLE and the variance by a weak inverse-gamma prior to prevent zero estimates caused by coarse time resolution. Abnormal cell displacement was assessed qualitatively, as positional defects in this context are typically unambiguous relative to the stereotypical embryonic arrangement. Chromatin and nuclear morphology defects were scored manually from time-lapse recordings using a binary scoring scheme for the presence or absence of two predefined, non-mutually exclusive categories: chromosome segregation defects (lagging chromosomes or chromatin bridges during mitosis), abnormal nuclear content (enlarged or multinucleated nuclei following division).

### Protein lysate preparation and western blot

For yeast samples, single colonies were streaked onto YPD/Bacto Agar plates and harvested after 3 days of incubation at 25°C or 37°C. Colonies were washed, snap-frozen in liquid nitrogen and resuspended in 0,2M NaOH. Following 10 minutes of incubation, samples were pelleted, resuspended in SDS loading buffer (50 mM TRIS pH6.8, 2% SDS, 10% glycerol, 0.1M DTT, 0.05% bromophenol blue) and boiled at 95°C for 5 minutes. For nematode samples, synchronized L1 larvae were incubated at 25°C for 40-48 h, producing a population enriched in gravid adults and early to mid-stage embryos. In case of heat shock induction samples, worms were then subjected to a 1h heat shock at 37 °C, followed by a 3h recovery at 25 °C before protein extraction.

Worms were washed off plates with M9, pelleted and resuspended in ice-cold lysis buffer: 50 mM HEPES pH 7.4, 150 mM NaCl, 2 mM MgCl_2_, 0.05% IGEPAL, and protease inhibitors (Roche, 11836153001). After snap-freezing, samples were lysed by sonication using a Bioruptor (UCD-200, Diagenode) for three cycles of 30 s ON/30 s OFF over 10 min at high energy in an ice-water bath, followed by centrifugation to collect the clarified supernatant. The protein concentration was quantified using a Bradford assay (Thermo Scientific, 23238). Samples were then mixed with SDS loading buffer, normalized (usually to 2 mg/mL) and boiled at 95°C for 5 min prior to loading. All samples were loaded onto NuPAGE Bis-Tris 4–12% gel (Invitrogen). After electrophoresis, samples were transferred to 0.45 µm polyvinylidene fluoride membrane (Thermo Scientific, 88518) and blocked with 5% non-fat milk in PBS-T with 0.1% of Tween20 (Sigma-Aldrich, P1379) for 1 h at room temperature. Following blocking, membranes were incubated overnight at 4 °C with primary antibodies diluted in blocking buffer: anti-KLMT-1 (clone 1A3-3E5; 1:60; Monoclonal Antibody Facility, Max Perutz Labs) or anti-PGK-1 (1:10,000; Invitrogen, 459250) as a yeast loading control. After incubation with the antibodies, membranes were washed with PBS-T, followed by incubation with anti-mouse HRP-conjugated secondary antibodies (1:10,000, Invitrogen, G-21040). Substrate detection was performed using ECL reagent (Cytiva, RPN2106) and imaged with ChemiDoc MP (Bio-Rad). An Alexa Fluor 647–conjugated anti-α-tubulin antibody (1:2000; Abcam, ab190573) was used as a loading control for worm lysates. Signal detection was performed as described above.Substrate detection was performed as described above.

### Preparation of embryos and adult worms for microscopy

For imaging of embryos, L4 larvae were selected one day prior to the experiment to ensure reproducibility and maximize egg yield. Young gravid adults were dissected with insulin syringe needles (29 G) in a drop of M9. If heat shock induction was required, embryos were first pipetted onto 5-cm modified NGM plates and incubated at 37°C for 1h. All embryos were then transferred in bulk into 8-well imaging chambers (Thermo Scientific, 155411) containing M9 medium and chambers were sealed with parafilm. For imaging of adult worms, the animals were anaesthetized using levamisole (1 mM in M9), transferred to microscope slides with agar pads and sealed with cover slips.

### Imaging of embryos and adult worms

DIC and fluorescence images of adult worms were acquired on an Axio Imager.Z2 (Zeiss) equipped with a 20×/0.8 Plan-Apochromat objective and a Hamamatsu Orca Flash 4 camera. The following filter settings were used: excitation 480/40 nm, emission 510LP. Images were stitched prior to export. Brightfield and fluorescence images of embryos were acquired on a spinning disc confocal microscope (Olympus IX83 inverted microscope with a Yokogawa W1 disc and Hamamatsu Orca Fusion camera). With the exception of fig. S2A (40X/0.75 objective), a 100X/1.4 oil immersion objective was used for all experiments. Laser power and exposure times were kept constant within experiments. For most imaging, settings were optimized to minimize phototoxicity (low laser power (10-15%), long exposure times (700-800 ms), coarse-grained imaging regime with 8 x 1 µm step size acquired every 30 minutes). For co-localization and foci size quantification experiments, we chose altered settings (40-80% laser power, 100 ms exposure, 50 x 0.1 µm step size) and a modified setup (dual Hamamatsu Orca cameras, piezoelectric Z stage) for maximized spatial and temporal resolution.

### Image post-processing and analysis

Representative micrographs were processed in Fiji. Unless stated otherwise, all images within a figure panel are shown at equal brightness/contrast settings. Background subtraction was performed by average intensity subtraction. Single-timepoint quantification and co-localization analysis of foci (KLMT-1, SAS-4, SPD-5) was performed using a custom Python package based on the PunctaFinder algorithm (*61*), which enables detection of foci in low signal-to-noise settings by combining global and local thresholding for segmentation. Parameters were chosen by benchmarking against manual foci picking and kept constant across all quantifications. Manual correction of total foci numbers was applied in rare cases where dim foci were not correctly identified. To assess the degree of spatial overlap between GFP and mScarlet foci, we calculated Manders colocalization coefficients for both fluorophores (M1, M2): 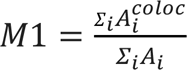 and *M*2 = 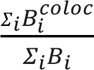, with A and B being pixel intensities of both fluorophore foci. Foci size was quantified in Fiji. Briefly, brightness and contrast settings were normalized, thresholding was applied (Otsu, range 180-65535) and particle size was measured (minimum size 0.05 µm^2^, circularity 0.2-1). Multi-timepoint foci counting was performed manually in Fiji. Brightness/contrast settings were normalized, and images were Z-projected (maximum intensity), background-subtracted and median-filtered (0.1 pixel size). The 8-cell stage was defined as timepoint 0. GFP and mScarlet foci were counted for every timepoint until comma stage using the multi-point selection tool. Only embryos that progressed normally through development were used for analysis. For all analyses described above, at least two individual imaging experiments were conducted. All imaging analysis raw data is displayed in Supplemental Data S4.

### De-novo protein structure and interaction prediction

All wild-type or mutant KLMT-1 / AIR-1 interactions were predicted with AlphaFold3 (*36*). Predictions were run at least twice with independent seeds. AlphaFold2 predictions of KLMT-1/AIR-1 were generated using the ColabFold implementation of AlphaFold2-multimer (version 1.3.0, MMseqs2 option) (*62*). PAE plots were generated using PAEViewer (*63*). The KLMT-1/AIR-1 interface was visualized using AlphaBridge (*64*) using default settings. Pairwise predictions of KLMT-1 and centrosomal proteins were also run in Alphafold3; candidates were selected from UniProt (*C. elegans* proteins annotated as centrosomal by GO:0005813). Protein IDs, input sequences and PTM/iPTM values of this dataset are listed in Supplemental Data S5.

### Statistical analysis & software

Representation and analysis of all genotyping, phenotyping, and fluorescence image quantifications was performed in GraphPad Prism10 (version 10.0.3). Figures and figure legends provide descriptions of the specific statistical tests used, error bars and sample sizes. For all tests, 0.05 was set as the significance threshold. If applicable, appropriate corrections for multiple comparisons were made. Multiple sequence alignments were generated using MUSCLE via the EMBL-EBI web server and alignments were visualized using Jalview (version 2.11.5.1). All structural models were generated in ChimeraX (version 1.10). Adobe Illustrator (version 30.3) was used for assembling figures and making illustrations.

