## Supplementary Materials for "Evolution of an Aurora Kinase A Inhibitor from an Essential tRNA Synthetase"

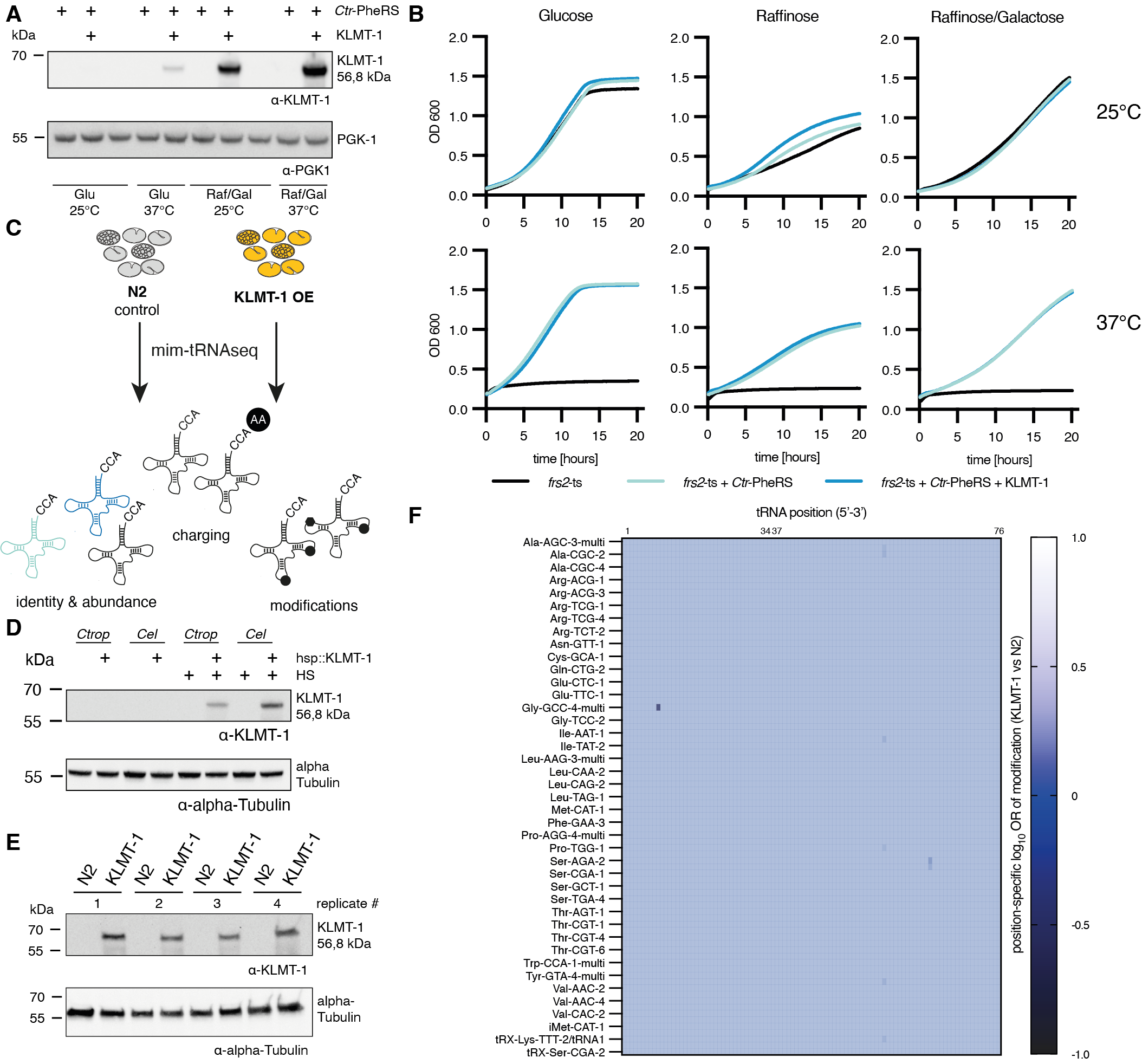

**Fig. S1. KLMT-1 does not hinder PheRS activity nor overall tRNA homeostasis. (A)** Western blot of lysates from *S. cerevisiae* *frs2*-ts cells transfected with *Ctr* PheRS and/or KLMT-1 expression constructs and cultured in different media and temperature conditions. KLMT-1 is induced in the presence of raffinose; weak leaky expression is observed in the presence of glucose at 37°C. (**B)** Liquid culture of *S. cerevisiae* frs2 (ts) co-expressing *C. tropicalis* PheRS +/- KLMT-1 at permissive (25°C) and restrictive (37°C) temperature in YPD with glucose, raffinose or raffinose/galactose; induction of PheRS expression by raffinose rescues growth, regardless of KLMT-1 co-expression. **C)** Schematic representation of the in- and output of mim-tRNAseq. **D)** Western blot of lysates from mixed-stage hsp::KLMT-1 *C. tropicalis* and *C. elegans* lines versus wild-type controls. Heat shock induces KLMT-1 expression in both species. **E)** Western blot of lysates from *C. elegans* embryos collected for tRNA sequencing. KLMT-1 is detected at comparable levels in all KLMT-1 OE samples. **F)** Heatmap illustrating the position-specific likelihood of a tRNA modification being enriched or depleted in KLMT-1-overexpressing *C. elegans* embryos over control. All distinguishable cytosolic tRNAs are shown; the anticodon loop residues 34/37, a hotspot of crucial modifications, is indicated above. Experiment in biological quadruplicates.

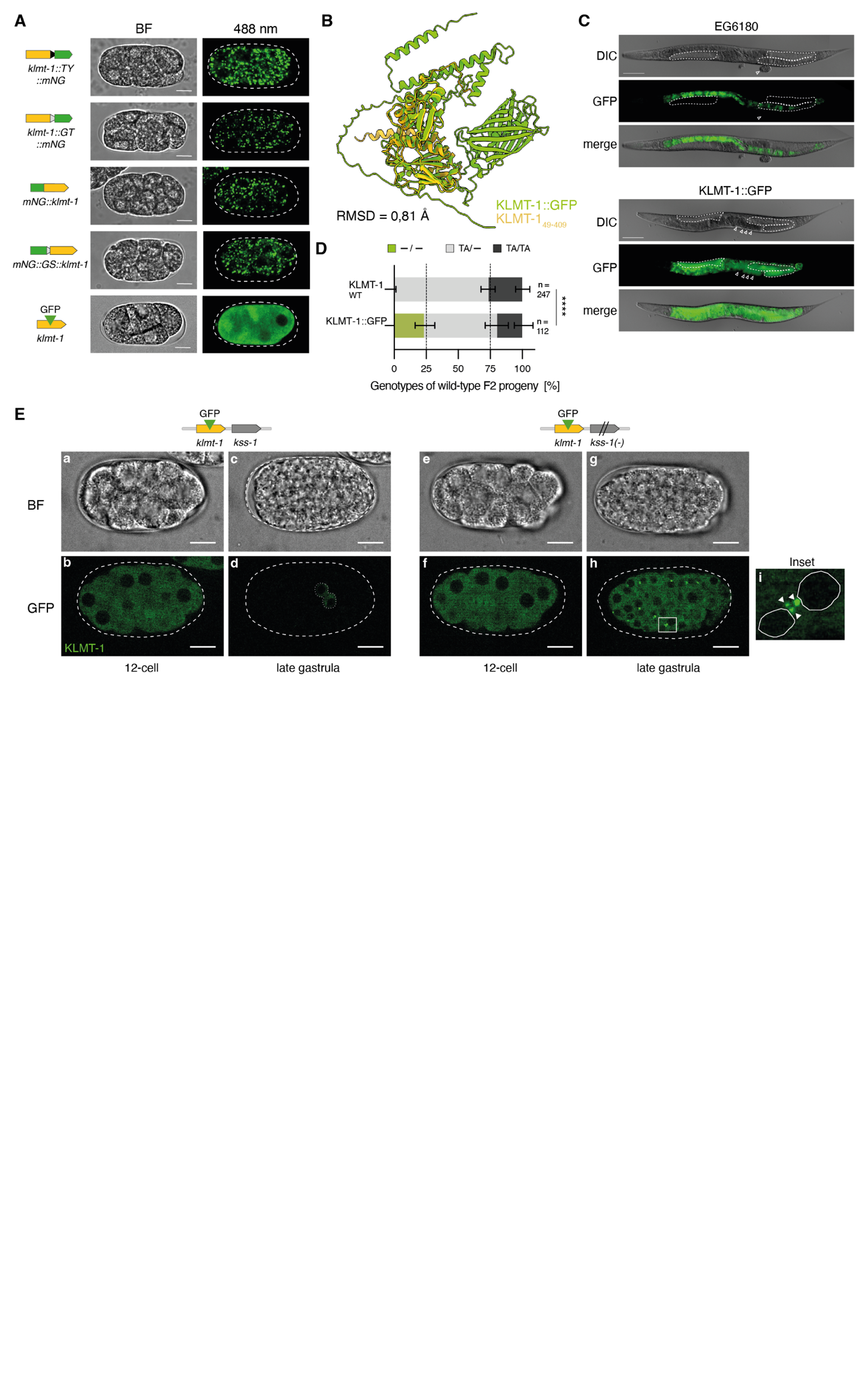

**Fig S2. An internal fluorescent tag prevents KLMT-1 aggregation in vivo. (A)** Brightfield and fluorescence micrographs of pre-gastrulation embryos expressing mEGFP- or mNeonGreen (mNG)- tagged variants of KLMT-1. All mNG-tagged constructs display cytoplasmic aggregation, while KLMT-1::GFP does not. Scale bar is 10 µm. Brightness and contrast were individually adjusted to best represent the intracellular distribution of KLMT-1. **(B)** AlphaFold 3 predictions of KLMT-1::GFP (green) aligned to the core region of KLMT-1 (AA:49-409, yellow), highlighting the conserved fold. RMSD (pruned) = 0,81 Angstroms. **(C)** DIC and fluorescence micrographs of wild-type (EG6180) and KLMT-1::GFP-expressing *C. tropicalis* adult hermaphrodites. Germline (dashed line) and embryos (arrowheads) are fluorescent only in the presence of KLMT-1::GFP. Scale bar is 100 µm. Autofluorescence in the gut is observed in all animals. **(D)** Genetic crosses showing the loss of toxicity of KLMT-1::GFP (*n*_(-/-)_ = 26/112, *P* < 0.0001, Fisher’s exact test). **(E)** Brightfield and fluorescence confocal micrographs of KLMT-1::GFP-expressing embryos in the presence (a-d) or absence (e-i) of KSS-1. KLMT-1 is present in all early embryos but efficiently degraded by KSS-1 (c,d). In the absence of KSS-1, KLMT-1 forms perinuclear foci (i) during gastrulation. Inset at 5X. Scale bar is 10 µm.

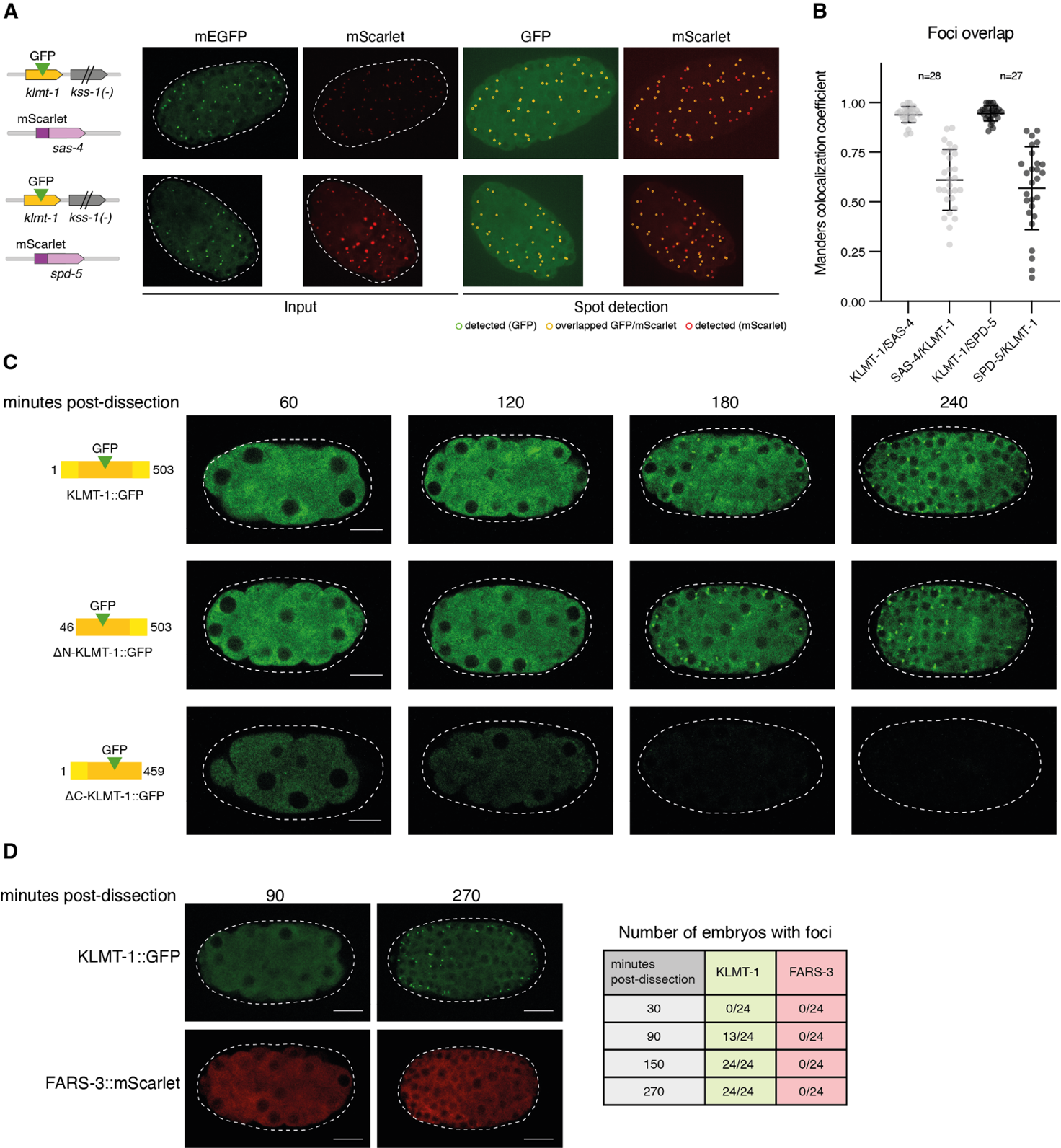

**Fig S3. KLMT-1, but not its ancestral paralog FARS-3, is dynamically localized to centrosomes. (A)** Illustration of the spot detection pipeline. Left: Z-projected input images from *C. tropicalis* embryos co-expressing KLMT-1::GFP with mScarlet-tagged SAS-4 and SPD-5, respectively. Right: spot detection algorithm recognizes foci in both channels and calculates Manders colocalization coefficients for every embryo. **(B)** Dot plot showing the degree of colocalization of KLMT-1::GFP foci with SAS-4 and SPD-5 foci, respectively. KLMT-1 strongly colocalizes with both proteins, SAS-4 and SPD-5 partially colocalize (to a similar extent) with KLMT-1. Mean +/- SD. Dots represent individual embryos from 2 experiments. **(C)** Fluorescence micrographs of *C. tropicalis* embryos expressing full-length or terminally truncated versions of KLMT-1::GFP over time. Foci are visible from 180 min onwards. N-terminal truncation does not impact fluorescence or foci formation, C-terminal truncation leads to lower initial levels and rapid degradation of KLMT-1. Scale bar is 10 µm. **(D)** Comparison of KLMT-1::GFP with mScarlet-tagged FARS-3. While all KLMT-1::GFP expressing embryos display foci formation (24/24 embryos at 270 minutes post-dissection), FARS-3 remains cytoplasmic (0/24).

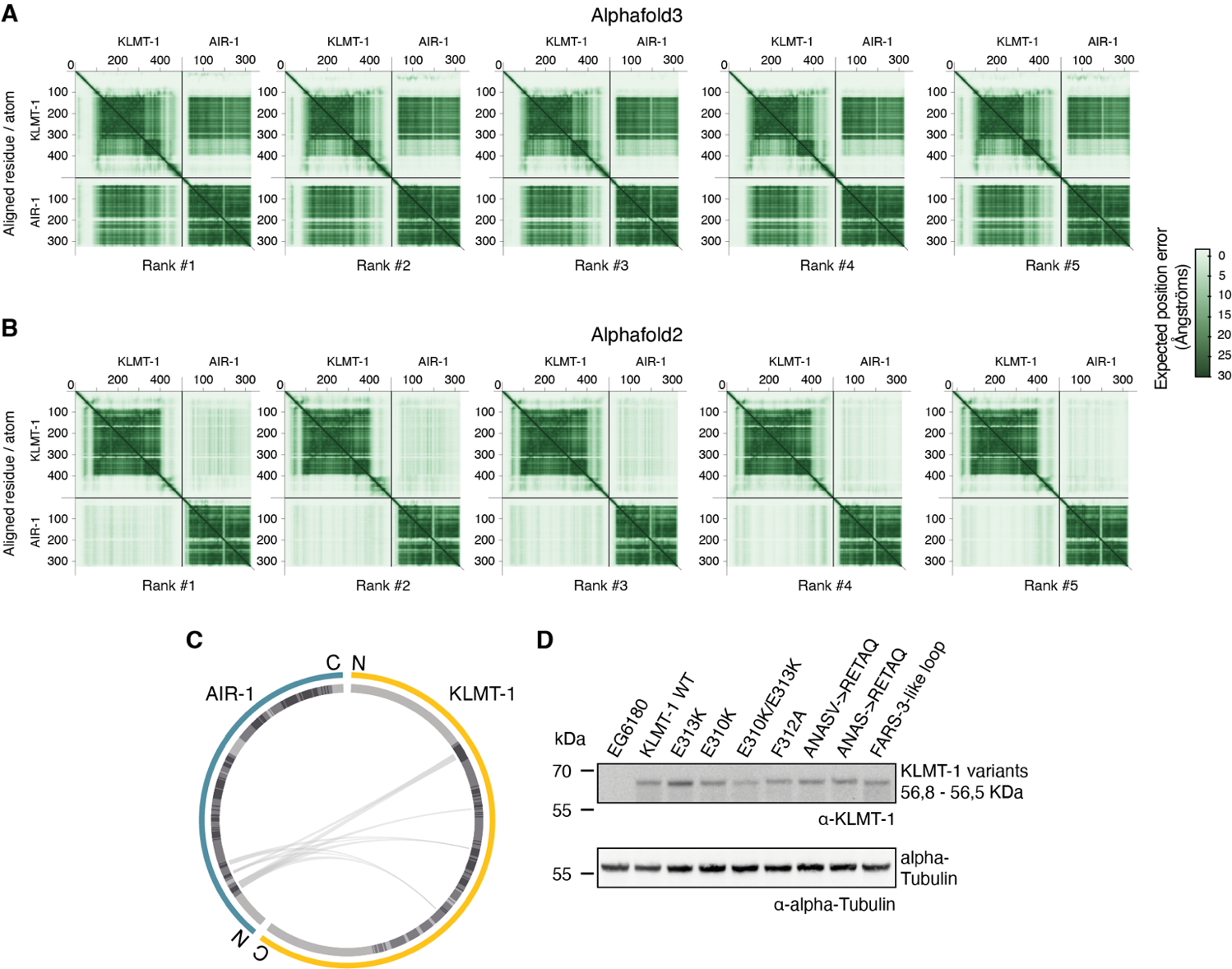

**Fig S4. AlphaFold2 and AlphaFold3 predictions of the KLMT-1–AIR-1 interaction, and expression levels of KLMT-1 mutants (A)** PAE plots for all AlphaFold3 models of the KLMT-/AIR-1 interaction. An interaction is confidently predicted in 5/5 models. **(B)** PAE plots for all AlphaFold2 models of the KLMT-/AIR-1 interaction. 0/5 models confidently predict an interaction. **(C)** AlphaBridge visualization of the predicted interface between KLMT-1(yellow) and AIR-1 (blue). Several regions of KLMT-1 contribute to the interface; interacting residues of AIR-1 are restricted to the N-terminal N-lobe. **(D)** Western blot comparing the expression levels of wild-type and mutant versions of KLMT-1 from the endogenous *klmt-1* locus generated using CRISPR/Cas editing.

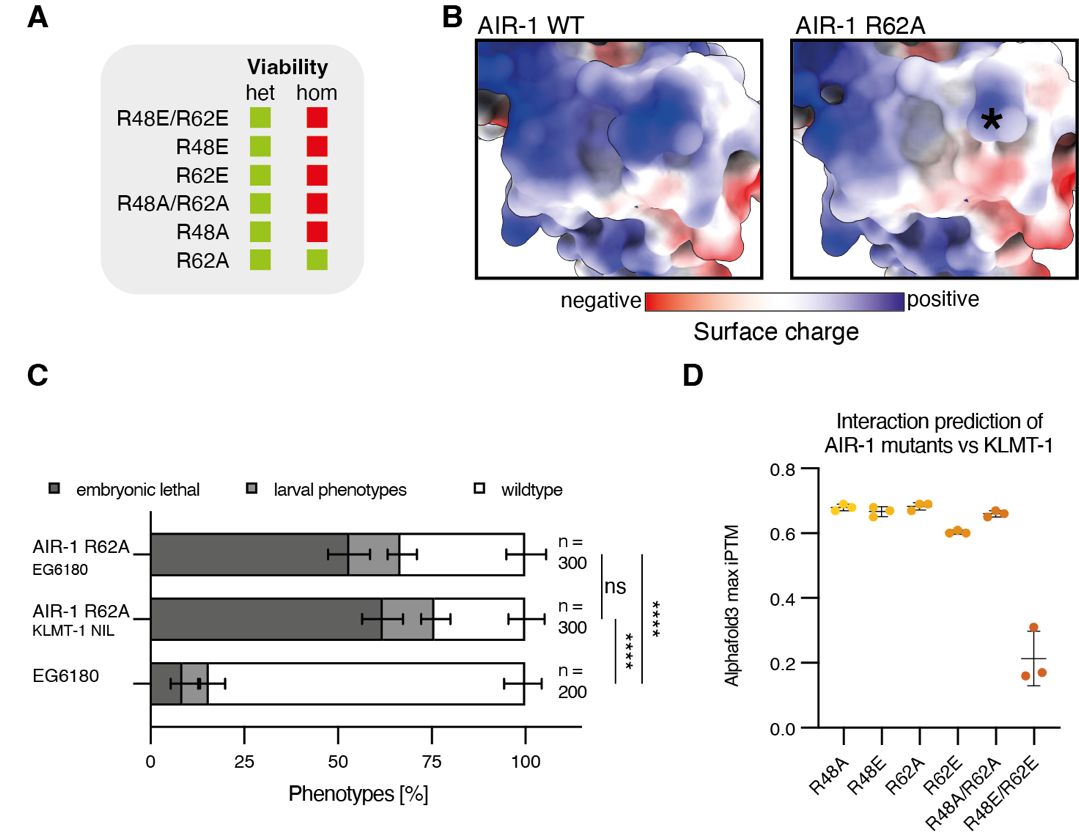

**Fig S5. Characterization of AIR-1 mutants in highly conserved residues predicted to interact with KLMT-1. (A)** Table summarizing the attempted CRISPR single and double edits of AIR-1 at the KLMT-1 interface. Only the R62A mutation yielded viable homozygous progeny. **(B)** The surface charge of AIR-1 at the N-lobe in wild-type versus R62A. The location of residue 62 is indicated by an asterisk. **(C)** Phenotyping of wild-type (EG6180) or AIR-1 R62A (EG6180 and KLMT-1 NIL background) *C. tropicalis* embryos. AIR-1 R62A causes severe pleiotropic defects (≥ 200/300 affected; *P*adj < 0.0001, Fisher's exact test with Bonferroni correction). **(D)** Effect of mutations at the AIR-1-KLMT-1 interface on interaction as predicted by AlphaFold3. Shown are maximum iPTM values of 3 independent predictions per pair. Only the R48E/R62E mutation is robustly predicted to disrupt binding.

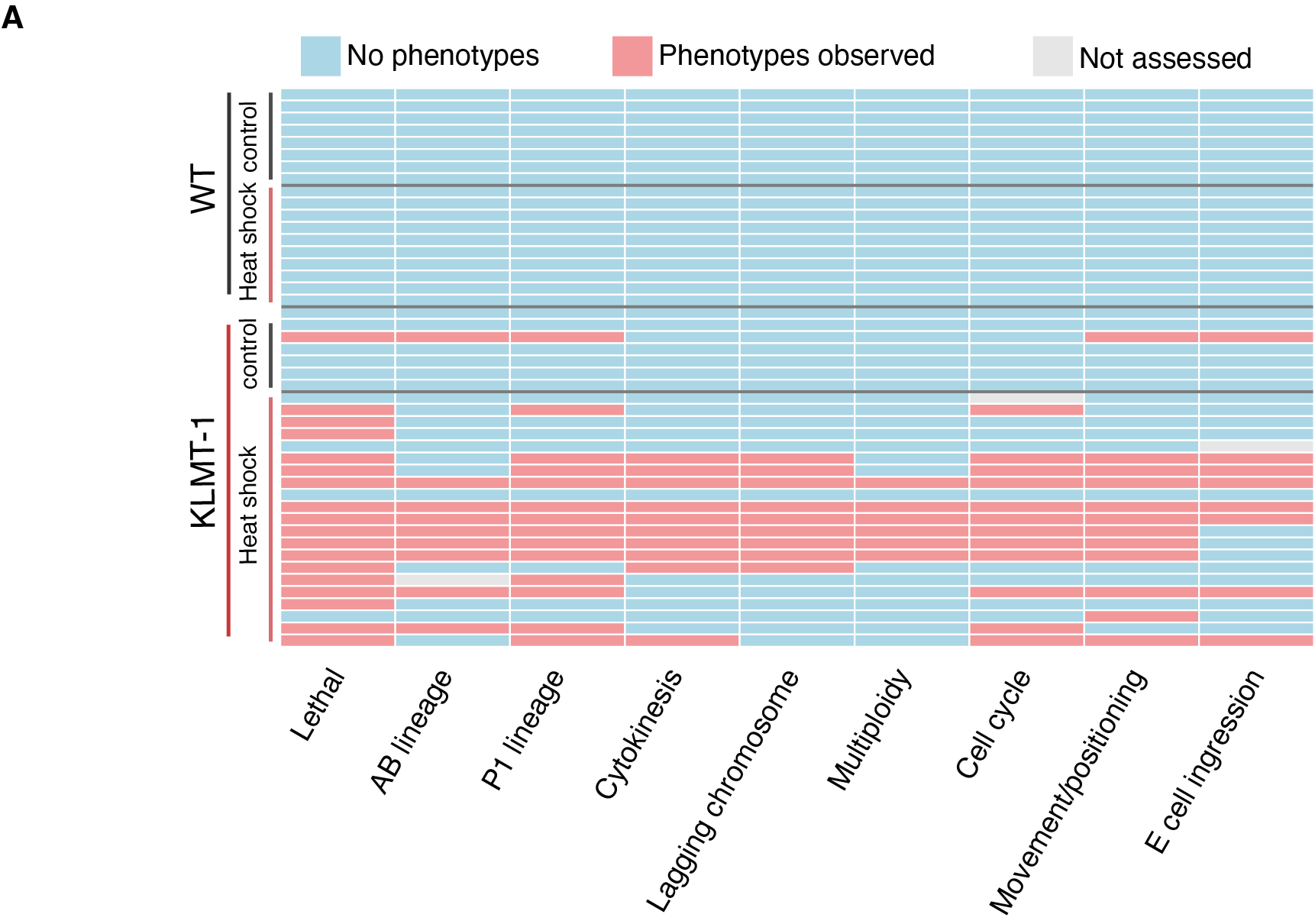

**Fig. S6. Quantification of the effect of KLMT-1 overexpression on *C. elegans* embryonic development.** Heatmap illustrating the diverse effects of KLMT-1 overexpression in *C. elegans* embryos. No phenotypes were observed in control embryos, independent of heat shock treatment. Each line represents an individual embryo.

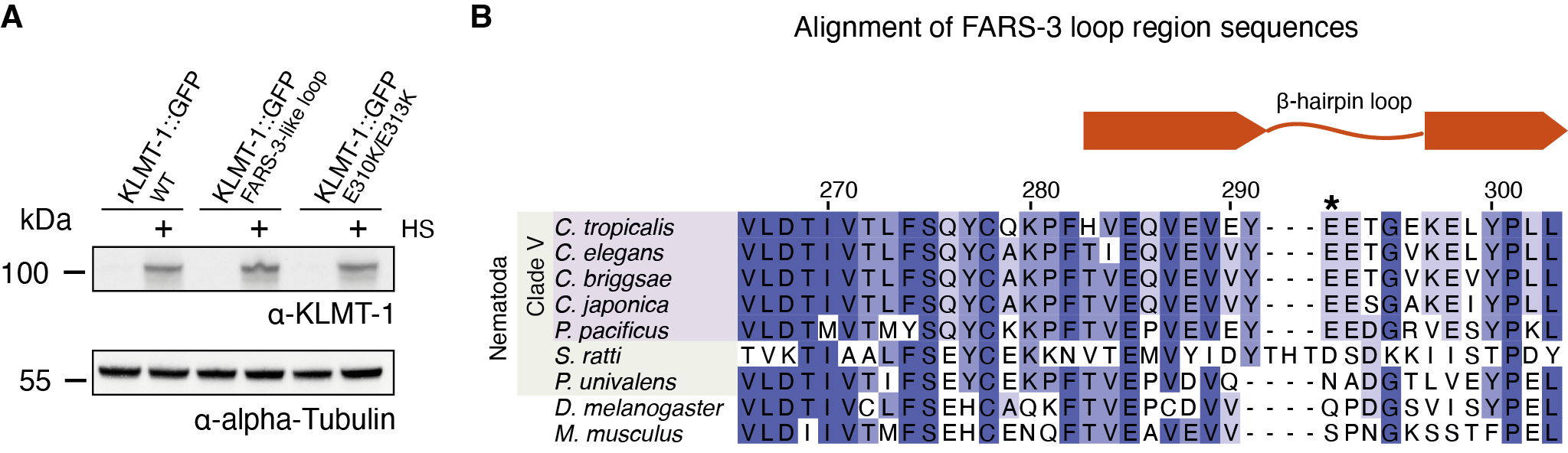

**Fig S7. Validation of KLMT-1 expression levels in different mutant lines and conservations of the FARS-3 beta hairpin loop. (A)** Western blot showing comparable expression of wild-type or loop mutant KLMT-1::GFP after heat-shock induction in *C. tropicalis*. **(B)** FARS-3 E292 is conserved among Clade V nematodes. Alignment of FARS-3 protein sequences from closely and distantly related species. *C. tropicalis* E^292^ is indicated by an asterisk. Sequence conservation displayed by shades of blue (darker = more conserved). The β-hairpin loop containing E^292^ is visualized above (orange; bars for beta sheets, line for connecting loop region).

Table S1.

List of *C. tropicalis* and *C. elegans* strains used in this study. Strains printed in bold were created for this work.

| ***C. tropicalis*** | | | | |
| --- | --- | --- | --- | --- |
| **Strain** | **Short name** | **Genotype** | **Description** | **Used in Figure** |
| EG6180 | EG6180 | *C. tropicalis* natural isolate (wild-type) | Wild isolate from El Yunque, Puerto Rico. Lat 18.3 Lon -65.8. Found in rotting fruit by M. Ailion and E. Jorgensen. Source: Christian Braendle | 1B/E, S1D, S2D, 3D/E, S3D, 4C, S4D,5B/F, S5C |
| QX2343 | KLMT-1 NIL | *qqIR47(V:1.3-1.8 Mb; NIC203 > EG6180); EG6180 Mito* | NIC203 TA element on Chr. V introgressed into EG6180 background. Ben-David et al (2021) | 1B, S2D, 3D/E, S4D, 4C, 5B |
| INK140 | KLMT-1::GT::mNG | *klmt-1(abu86[klmt-1::GT::mNeonGreen])* V*; qqIR47* | C-terminal mNeonGreen tagged KLMT-1 (with GT linker), QX2343 background | S2A |
| INK169 | KLMT-1::TY::mNG | *klmt-1(abu107[klmt-1::2xTy1::mNeonGreen])* V*; qqIR47* | C-terminal mNeonGreen tagged KLMT-1 (with 2xTy1 linker), QX2343 background | S2A |
| INK184 | mNG::KLMT-1 | *klmt-1(abu110[mNeonGreen::klmt-1])* V; *qqIR47* | N-terminal mNeonGreen tagged KLMT-1, QX2343 background | S2A |
| INK217 | mNG::GS::KLMT-1 | *klmt-1(abu128[mNeonGreen::GS::klmt-1])* V; *qqIR47* | N-terminal mNeonGreen tagged KLMT-1 (with GS linker), QX2343 background | S2A |
| INK285 | Chr. IV SLP | *abuSi9[dpy-10 + sup-35 gRNA targets::(HygR(aa52-341)::rps-20 3' UTR::LoxP, IV:6.9Mb]* IV | Chr. IV synthetic landing pad with split hygromycin resistance in EG6180 background. Only allows for somatic transgene expression. Parental line for all *C. tropicalis* SLP lines. |  |
| INK318 | Heat-shock inducible KLMT-1 | *klmt-1(abuSi19[hsp-16.11p::klmt-1::tbb-2 3' UTR + HygR(+); abuSi9])* IV | Heat shock inducible expression of KLMT-1. Expressed from Chr. IV synthetic landing pad (INK285). Only somatic expression | 1E, S1D |
| **INK873** | Heat-shock inducible KLMT-1::GFP | *abuSi30[hsp-16.11p::KLMT-1[Q234_ins_mEGFP]::tbb-2 3' UTR + HygR(+); abuSi9]* IV | Heat shock inducible expression of internally mEGFP tagged KLMT-1. Expressed from Chr. IV synthetic landing pad (INK285). Only somatic expression | S2D, 5C, S7A |
| **INK966** | KLMT-1::GFP + KSS-1 | *klmt-1(abu504[Q234_ins_mEGFP])* V; *qqIR47* | Internally mEGFP tagged KLMT-1 (at position 234 with flanking 5xG linkers), QX2343 background | S2D/E |
| **INK1021** | FARS-3::mScarlet | *fars-3(abu397[FARS-3::mScarlet])* II*; mEGFP(abuSi116[rpl-36p::SV40::mEGFP::egl-13-NLS::rpl36-3' UTR + HygR(+), abuSi33])* I | FARS-3 tagged with mScarlet on the C-terminus, additional NLS-mEGFP, EG6180 background | S3D |
| **INK1041** | KLMT-1::GFP | *klmt-1(abu504[Q234_ins_mEGFP])* V; *kss-1*(abu506[p.Ile6ThrfsTer50]) V; *qqIR47* | Internally mEGFP tagged KLMT-1 with a *kss-1* knockout mutation. QX2343 background | S2E |
| **INK1184** | Heat-shock inducible KLMT-1::GFP E310K/E313K | *abuSi151[hsp-16.11p::KLMT-1[Q234_ins_mEGFP ][p.E310K; p.E313K)]::tbb-2 3' UTR + HygR(+); abuSi9]* IV | Heat shock inducible expression of internally mEGFP tagged mutant KLMT-1 (E310K/E313K). Expressed from Chr. IV synthetic landing pad (INK285). Only somatic expression | 5C, 7A |
| **INK1190** | Heat-shock inducible KLMT-1::GFP FARS-3-like loop | *abuSi157[hsp-16.11p::KLMT-1[Q234_ins_mEGFP]* [p.P315K/delRAL(316-318)]::*tbb-2 3' UTR + HygR(+); abuSi9]* IV | Heat shock inducible expression of internally mEGFP tagged mutant KLMT-1 (deletion of AA 316-318, P315K). Expressed from Chr. IV synthetic landing pad (INK285). Only somatic expression. | 5C, S7A |
| **INK1225** | KLMT-1::GFP E310K/E313K | *klmt-1(abu436[p.E310K; p.E313K])* V; *qqIR47* | KLMT-1 charge-reversing mutant (E310K/E313K), QX2343 background | 3D, S4D |
| **INK1230** | KLMT-1::GFP FARS-3-like loop | *klmt-1(abu441[p.316-318del; p.P315K])* V; *qqIR47* | KLMT-1 FARS-3 like loop mutant (deletion of AA 316-318, P315K), QX2343 background | 5B, S4D |
| **INK1292** | KLMT-1::GFP and mScarlet-SAS-4 | *sas-4 (abu198[mScarlet-I3::sas-4])* III, *klmt-1(abu504[Q234_ins_mEGFP])* V*; kss-1(abu506[p.Ile6ThrfsTer50])* V; *qqIR47* | N-terminal mScarlet-I3 tagged SAS-4, INK1041 background | 2A-D, S2C, S3A-D |
| **INK1325** | KLMT-1 NIL AIR-1 R62A | *air-1(abu578[p.R62A])* V; *qqIR47* | AIR-1 point mutation (R62A), QX2343 background | 4C, S5C |
| **INK1328** | EG6180 AIR-1 R62A | *air-1(abu581[p.R62A])* V | AIR-1 point mutation (R62A), EG6180 background | 4C, S5C |
| **INK1376** | KLMT-1 FARS-3-like helix I | *klmt-1(abu282*[KLMT-1 p.ANAS(132-135)RETAQ]*)* V; *qqIR47* | KLMT-1 with FARS-3 like helix in QX2343 background | 3E, S4D |
| **INK1396** | KLMT-1::GFP and mScarlet-SPD-5 | *spd-5 (abu599[mScarlet-I3::spd-5])* I; *klmt-1(abu504[Q234_ins_mEGFP])* V; *kss-1(abu506[p.Ile6ThrfsTer50])* V | N-terminal mScarlet-I3 tagged SPD-5, INK1041 background | 2A/B, S3A/B |
| **INK1527** | FARS-3 E292A | *fars-3(abu649[p.E292A])* II | FARS-3 point mutation (E292A), EG6180 background | 5F |
| **INK1530** | FARS-3 E292K | *fars-3(abu649[p.E292K])* II | FARS-3 point mutation (E292K), EG6180 background | 5F |
| **INK1533** | KLMT-1 E313K | *klmt-1(abu655[p.E313K])* V; *qqIR47* | KLMT-1 point mutation (E313K), QX2343 background | 3D, S4D |
| **INK1574** | KLMT-1 E310K | *klmt-1(abu664[p.E310K])* V; *qqIR47* | KLMT-1 point mutation (E310K), QX2343 background | 3D, S4D |
| **INK1601** | KLMT-1 NIL AIR-1 A62R | *air-1(abu676[p.A62R])* V | AIR-1 point mutation (A62R), INK1325 background. Restores AIR-1 wild-type sequence. | 4C |
| **INK1690** | KLMT-1 FARS-3-like helix II | *klmt-1(abu696[p.ANASV(132-136)RETAQ])* V | KLMT-1 with FARS-3 like helix in QX2343 background (alternative design) | 5B, S4D |
| **INK1717** | KLMT-1::GFP ∆N and mScarlet::SAS-4 | *sas-4 (abu198[mScarlet-I3::sas-4])* III; *kss-1(abu506[p.Ile6ThrfsTer50])* V; *klmt-1(abu721[Q234_ins_mEGFP][p.2-45del])* V | Internally mEGFP tagged KLMT-1 with N-terminal deletion (AA 2-45), KSS-1 knockout and mScarlet::SAS-4. | S3C |
| **INK1770** | KLMT-1::GFP ∆C and mScarlet::SAS-4 | *sas-4 (abu198[mScarlet-I3::sas-4])* III; *kss-1(abu506[p.Ile6ThrfsTer50])* V; *klmt-1(abu729[Q234_ins_mEGFP][p.460-503del])* V | Internally mEGFP tagged KLMT-1 with C-terminal deletion (AA 460-503). KSS-1 knockout and mScarlet::SAS-4. | S3C |

| ***C. elegans*** | | | | |
| --- | --- | --- | --- | --- |
| **Strain** | **Short name** | **Genotype** | **Description** | **Used in Figure** |
| N2 | N2 | *C. elegans* reference strain (wild-type) | Caenorhabditis elegans reference strain. Isolated from mushroom compost near Bristol, England by L.N. Staniland. | 1E-G, S1D-F |
| RW10226 | Lineage tracing | *ItIs37[pie-1p::mCherry::his-58 + unc-119(+)]* IV*. stIs10226 [his-72p::HIS-24::mCherry::let-858 3' UTR + unc-119(+)]* IV | Ubiquitous histone-cherry expression in C. elegans suitable for lineage analysis. Created in the Waterston lab. Source: CGC | 4E/F, S6 |
| INK252 | Heat-shock inducible KLMT-1 | *abu157[hsp-16.41p::(Frt)::klmt-1::unc-54 3'UTR]* II | Heat shock inducible expression of KLMT-1 from Chr. II, N2 background. | 1E-G, S1D-F |
| INK800 | Heat-shock inducible KLMT-1 (lineage tracing) | *abu157[hsp-16.41p::(Frt)::klmt-1::unc-54 3'UTR]* II; *ItIs37 [pie-1p::mCherry::his-58 + unc-119(+)]* IV. *stIs10226 [his-72p::HIS-24::mCherry::let-858 3' UTR + unc-119(+)]* IV | Heat shock inducible expression of KLMT-1 from Chr. II in RW10226 background for lineaging analysis. | 4E/F, S6 |

Table S2.

List of repair templates and primers for *C. tropicalis* Chr. IV synthetic landing pad injections. Amplicon size = insertion size + 2,2 kb (empty landing pad).

| Lines generated | Insertion (short name) | Transgene inserted | Repair template sequence (plasmid) | Genotyping primer - 1 | Genotyping primer - 2 | Insertion size [bp] |
| --- | --- | --- | --- | --- | --- | --- |
| INK873 | heat-shock inducible (hsp) KLMT-1::GFP | *abuSi30[hsp-16.11p::klmt-1(1-234)::mEGFP::klmt-1(235-503)::tbb-2 3' UTR + HygR(+)] IV* | pAB0383 | TCCAATCTCGCTCTTCAACTCGT | TGTTCGCCGTACAGAGAACATCT | 4346 |
| INK1184 | hsp-KLMT-1 (E310K/E313K) | *abuSi151[hsp-16.11p::klmt-1(1-234)::mEGFP::klmt-1(235-503 p.E310K/E313K)::tbb-2 3' UTR + HygR(+)] IV* | pAB0447 | TCCAATCTCGCTCTTCAACTCGT | TGTTCGCCGTACAGAGAACATCT | 4346 |
| INK1190 | hsp-KLMT-1 (FARS-3-like loop) | *abuSi157 [hsp-16.11p::klmt-1(1-234)::mEGFP::klmt-1(235-503 p.P315K del.316-318)::tbb-2 3' UTR + HygR(+)] IV* | pAB0449 | TCCAATCTCGCTCTTCAACTCGT | TGTTCGCCGTACAGAGAACATCT | 4337 |

Table S3.

List of guide RNAs, homology-directed repair templates and genotyping primers used for editing endogenous genes in this study.

| **Strain** | **Modification** | **gRNA sequence** | **Repair template sequence** | **Genotyping primer 1** | **Genotyping primer 2** | **Amplicon size [bp]** |
| --- | --- | --- | --- | --- | --- | --- |
| INK140 | *klmt-1::GT::mNG* | GAATGTATTTAACTTTCCAT | ccaagaactccttttcgacctcaaacacgtcacctttgatttgaagaatgaaaaaccaactcctagaattttcatcttgagaaaatttggtttgaaaataaaatacggatttcaggagccaccaaacatcacaaattttaatcggatctcctcaacgctttcgaagcttctggaactcgatgaaactaatccggacagtgtcttctcgagaaaacactcttccgaactgaagaaagtgactgccaaagtgctcaccctaatgaaaggagaagtcaaaagagctgagaagaatttgttaatgattcagagcaaagttggattggagatgaggtattttcgcacttcgcaacaaaacagcttcattttcgattttttcagtgacagtgagacggagaaagccgccgacgaacaatccgtggagaaaaagagaaactcctcgtcacagaatccaatggaaagtggtaccgtctccaagggagaggaggacaacatggcctccctcccagccacccacgagctccacatcttcggatccatcaacggagtcgacttcgacatggtcggacaaggaaccggaaacccaaacgacggatacgaggagctcaacctcaagtccaccaaggtaagtttaaacatatatatactaactaaccctgattatttaaattttcagggagacctccaattctccccatggatcctcgtcccacacatcggatacggattccaccaatacctcccatacccagacggaatgtccccattccaagccgccatggtcgacggatccggataccaagtccaccgtaccatgcaattcgaggacggagcctccctcaccgtcaactaccgttacacctacgagggatcccacatcaaggtaagtttaaacagttcggtactaactaaccatacatatttaaattttcagggagaggcccaagtcaagggaaccggattcccagccgacggaccagtcatgaccaactccctcaccgccgccgactggtgccgttccaagaagacctacccaaacgacaaggtaagtttaaacatgattttactaactaactaatctgatttaaattttcagaccatcatctccaccttcaagtggtcctacaccaccggaaacggaaagcgttaccgttccaccgcccgtaccacctacaccttcgccaagccaatggccgccaactacctcaagaaccaaccaatgtacgtcttccgtaagaccgagctcaagcactccaagaccgagctcaacttcaaggagtggcaaaaggccttcaccgacgtcatgggaatggacgagctctacaagtaaatacattctttcccccctttccccgcttcatcaatgtttcgaagaaaacgtttcattgtgttgtttgtgatgagatttggtttgtttttcagtgccactctttcagtggcaagtatataaattgtaatgtttttatttaaaaattatctctatatttatatttgccaaaaaatctggagtttaactgaaaaaacgcgcgtaaactccagatttttcttttttttttgacctgaaaatttaatttgccgcctaagccccccttcccctcacttttctcgattttcccccatttcccctttgttcccactcaattttcctgattcgaaaccctaatccgaaatagcacggggattacgttatcgagtctcttttttgaactcggcgcatagagctcagttcaaaaattatgccatgttttggataactttgcgttttttgattcccccaccctttcatcgtcatcccgtcatttttccagacgcggcgcaacgatttgagagtattgagcgagaaaaggaagaaaaatgctccgggaatcccaatggcttccgtggggcgtggtcaaaaaccacccgaaatgagct | TCCTCGAAGTGATGATACCGCC | GCTCAGAGGCTGATTTCCGAGT | 2311 |
| INK184 | *mNG::klmt-1* | CTGATTTTCGGACATTTCTC | gttccgaaaaattttctccgcttgttttcgctcttatttttaaacatttcttgtttcttagcatgcttttgcaattttgtaaaaattattcagcgtcaaaaatcagaaaaataagacgaaacaatgacttcgaaccaaactgttataattattaaaattttaattgaagaggaacaatattatacaagaaaaattgataaattcgtcaaaaataactagatcgtgtcgagacccgatggacaattaatcgcatgcgccttaaataccgtatactgaggaataaatttattcaccttattcttcgctaaatctttcgaatttttttgttttttccgtggtttttctttgctattttcgaatttaaattaaaaatacggctaataatctcaatattccagagaaatggtctccaagggagaggaggacaacatggcctccctcccagccacccacgagctccacatcttcggatccatcaacggagtcgacttcgacatggtcggacaaggaaccggaaacccaaacgacggatacgaggagctcaacctcaagtccaccaaggtaagtttaaacatatatatactaactaaccctgattatttaaattttcagggagacctccaattctccccatggatcctcgtcccacacatcggatacggattccaccaatacctcccatacccagacggaatgtccccattccaagccgccatggtcgacggatccggataccaagtccaccgtaccatgcaattcgaggacggagcctccctcaccgtcaactaccgttacacctacgagggatcccacatcaaggtaagtttaaacagttcggtactaactaaccatacatatttaaattttcagggagaggcccaagtcaagggaaccggattcccagccgacggaccagtcatgaccaactccctcaccgccgccgactggtgccgttccaagaagacctacccaaacgacaaggtaagtttaaacatgattttactaactaactaatctgatttaaattttcagaccatcatctccaccttcaagtggtcctacaccaccggaaacggaaagcgttaccgttccaccgcccgtaccacctacaccttcgccaagccaatggccgccaactacctcaagaaccaaccaatgtacgtcttccgtaagaccgagctcaagcactccaagaccgagctcaacttcaaggagtggcaaaaggccttcaccgacgtcatgggaatggacgagctctacaagctttccgaaaatcagcgactttcgaacaacggttctgtagaagaattttatacggtaaaaatggtttttttaaaaagacaaagtgcttttttgagatagttctaagcggttttatgcgaaaacataatatttgcggcgattattaggttgaaaaaacaaattttccatggaaaaatgctattttcaagaaaaaaaatttgtttcttcttttccagcgaataccaacagaaaagtagcaattttaatggcgaaaaccattattttctataatttttcacctgaaaatattaaattgtcccgtttgctaacccattttgcatttgaaaaacataaaatcccattcgaaatctcaaaaatatttattttcaggctccggacgaatacgaaaagacaaccactgcatctaagcagtt | GCGAGGCCGTATGATCTCTCAT | TGGAGGAAGGTCTGGAGCATTG | 1752 |
| INK217 | *mNG::GS::klmt-1* | CTTCACCGACGTCATGGGAA | gagctcaacttcaaggagtggcaaaaggccttcaccgacgtgatgggtatggatgaactttacaagggtggaggtggatcaggaggtggaggttcaggtggaggtggatctctttccgaaaatcagcgactttcgaacaacggttctgtagaagaattttatacggtaaaaatggtttttttaaaaagaca | CTACACCTTCGCCAAGCCAATG | TTTTCGTATTCGTCCGGAGCCT | 591 |
| NA | *klmt-1::GFP* partial integration | CTTCAACTCATTCAAAATCTT,  GAAGATTTTGAATGAGTTGA | aatgcatcggtcttccacccatatttatccggtgcgattgttcgccagatttcattagatttggaatctctcagatttttgaatggattggtagctgataaccaggagacgggcaagaaaagaatgcaatatttcatcacagcacaggatttggacaaaatgagtggaccgtttgaatatcgggccgaagcttcaaacgtaatcaaattccgtccgttaaatcagacgaaggagcacacggcggatgagctgatgacgttgtactcatctgacagccatatgacagatttccttcaactcattcaaggtggaggtggaggtgtgtccaagggagaggaactcttcaccggagtcgtcccaatcctcgtcgagctcgacggagacgtcaacggacacaagttctccgtctcaggagagggagagggagacgccacctacggaaagctcaccctcaagttcatctgcaccaccggaaagctcccagtcccatggccaaccctcgtcaccaccctgacttacggagtccaatgcttctcccgttacccagaccacatgaagcagcacgacttcttcaagtccgccatgccagagggatacgtccaagagcgtaccatcttcttcaaggtaagtttaaacattaattaatactaactaaccctgattatttaaattttcaggacgacggaaactacaagacccgtgccgaggtcaagttcgagggagacaccctcgtcaaccgtatcgagctcaaggtaagtttaaacagttcggtactaactaaccatacatatttaaattttcagggaatcgacttcaaggaggacggaaacatcctcggacacaagctcgaatacaactacaactcccacaacgtctacatcatggccgacaagcaaaagaacggaatcaaggtcaacttcaaggtaagtttaaacatgattttactaactaactaatctgatttaaattttcagatccgtcacaacatcgaggacggatctgtccaactcgccgaccactaccaacaaaacaccccaatcggagacggaccagtcctcctcccagacaaccactacctctccacccaatccaagctctccaaggacccaaacgagaagcgtgaccacatggtcctcctggagttcgtcaccgctgccggaatcaccctcggaatggacgagctctacaagggtggaggtggaggtaatcttcctgtctacccggtaatcgctgataagaatggactgatatccttggtgccagtagtcgatgcggaacctatgaagatatcaatcgagacgaagagtttgatgattatcgtaacatcagttgataaagagaacggcatccgaatcctcaacaacgtcctcgccgctatttcgattcgaattgaaaaaccacttgtcgttgagcccgttttgattgaatacgagaaattcgaggaaccgagagccttagag | TGGACAAGAAGCACGGAGAGAC | GGCGGTATCATCACTTCGAGGA | 1470 |
| INK966 | *klmt-1::GFP* | CAACTACAACTCCCACTACC | gttcatctgcaccaccggaaagctcccagtcccatggccaaccctcgtcaccaccctgacttacggagtccaatgcttctcccgttacccagaccacatgaagcagcacgacttcttcaagtccgccatgccagagggatacgtccaagagcgtaccatcttcttcaaggtaagtttaaacattaattaatactaactaaccctgattatttaaattttcaggacgacggaaactacaagacccgtgccgaggtcaagttcgagggagacaccctcgtcaaccgtatcgagctcaaggtaagtttaaacagttcggtactaactaaccatacatatttaaattttcagggaatcgacttcaaggaggacggaaacatcctcggacacaagctcgaatacaactacaactcccacaacgtctacatcatggccgacaagcaaaagaacggaatcaaggtcaacttcaaggtaagtttaaacatgattttactaactaactaatctgatttaaattttcagatccgtcacaacatcgaggacggatctgtccaactcgccgaccactaccaacaaaacaccccaatcggagacggaccagtcctcctcccagacaaccactacctctccacccaatccaagctctccaaggacccaaacgagaagcgtgaccacatggtcctcctggagttcgtcaccgctgccggaatcaccctcggaatggacgagctctacaagggtggaggtggaggtaatcttcctgtctacccggtaatcgctgataagaatggactgatatccttggtgccagtagtcgatgcggaacctatgaagatatcaatcgagacgaagagtttgatgattatcgtaacatcagttgataaagagaacggcatccgaatcctcaacaacgtcctc | TGGACAAGAAGCACGGAGAGAC | GGCGGTATCATCACTTCGAGGA | 1818 |
| INK1021  (INK722) | *fars-3::mScarlet* | ATAACAATAATTTAGAGGAA | cccttccatgtggagccgtggaattcaacgtggaaccgttcctcggaggtggatcaatggtctcgaaaggagaggccgtcatcaaggagttcatgcgtttcaaggtccacatggagggatccatgaacggacacgagttcgagatcgagggagagggagagggacgtccatacgagggaacccaaaccgccaagctcaaggtcaccaaggtaagtttaaacatatatatactaactaaccctgattatttaaattttcagggaggaccactcccattctcctgggacatcctctccccacaattcatgtacggatcccgtgccttcatcaagcacccagccgacatcccagactactacaagcaatccttcccagagggattcaagtgggagcgtgtcatgaacttcgaggacggaggagccgtcaccgtcacccaagacacctccctcgaggacggaaccctcatctacaaggtaagtttaaacagttcggtactaactaaccatacatatttaaattttcaggtcaagctccgtggaaccaacttcccaccagacggaccagtcatgcaaaagaagaccatgggatgggaggcctccaccgagcgtctctacccagaggacggagtcctcaagggagacatcaagatggccctccgtctcaaggacggaggacgttacctcgccgacttcaaggtaagtttaaacatgattttactaactaactaatctgatttaaattttcagaccacctacaaggccaagaagccagtccaaatgccaggagcctacaacgtcgaccgtaagctcgacatcacctcccacaacgaggactacaccgtcgtcgagcaatacgagcgttccgagggacgtcactccaccggaggaatggacgaactctacaagtaaattattgttattgaatgttttctatggaattgttgtttttaaatttaaatattc | ATTCCATTGATCTCACCCCCTGG | TTCTGGCTGGGAAAAAGGGTTCT | 1506 |
| INK1041 | *kss-1* frameshift in  *klmt-1::GFP* | CTTATTCGACTCCCAGACGT | --- | TATCCCATCTGCCACGTGTTGA | TACCGATGAAAGAGGTGTCGCC | 349 |
| INK1225 | *klmt-1::GFP* E310K/E313K | GATTGAATACGAGAAATTCG | cgaattgaaaaaccacttgtcgttgagcccgttttgattgaatacaaaaagtttaaggaaccgagagccttagagctctccccaccgctctcttatcgc | ACGTTGTACTCATCTGACAGCCA | TTTTTGTAGGCACACGCGACTC | 539 |
| INK1230 | *klmt-1::GFP* FARS-3-like loop | AGAGCCTTAGAGCTCTCCCC | gtcgttgagcccgttttgattgaatacgagaaattcgaggaaaaagagctctccccaccgctctcttatcgcaagatgacagtgacg | ACGTTGTACTCATCTGACAGCCA | TTTTTGTAGGCACACGCGACTC | 530 |
| INK1292 | *klmt-1::GFP/mScarlet-sas-4* | CTTCGGATGACATTCTGAAC | tatttcatgtctattctttgcgagatgttttttctatacacctgttcagaatggactccaccgaggccgtcatcaaggagttcatgcgtttcaaggtccacatggagggatccatgaacggacacgagttcgagatcgagggagagggagagggacgtccatacgagggaacccaaaccgccaagctcaaggtcaccaaggtaagtttaaacatatatatactaactaaccctgattatttaaattttcagggaggaccactcccattctcctgggacatcctctccccacaattcatgtacggatcccgtgccttcatcaagcacccagccgacatcccagactactggaagcaatccttcccagagggattcaagtgggagcgtgtcatgatcttcgaggacggaggaaccgtctccgtcacccaagacacctccctcgaggacggaaccctcatctacaaggtcaagctccgtggaggaaacttcccaccagacggaccagtcatgcaaaagcgtaccatgggatgggaggcctccaccgagcgtctctacccagaggacgtcgtcctcaaggtaagtttaaacagttcggtactaactaaccatacatatttaaattttcagggagacatcaagatggccctccgtctcaaggacggaggacgttacctcgccgacttcaagaccacctacaaggccaagaagccagtccaaatgccaggagccttcaacatcgaccgtaagctcgacatcacctcccacaacgaggactacaccgtcgtcgagcaatacgagcgttccgtcgcccgtcactccaccggaggatcaggaggatcaggttcatccgaagacaacgagaacgatggaagaactcggaagccga | TGAATCAAGCTCTAGCAAAGCGG | ATCTGGGAAGACTGATGGCAGG | 1310 |
| INK1325 | KLMT-1 NIL *air-1* R62A | CGAGAAGAAAACGAGACGAA | ctggagtctcgacgatttcgacgttggcgctccacttggaaagggaaagtttggaaatgtgttcatcagtgccgagaagaaaacgagacgaattatcgctctgaaagttctcttcaaaacgc | TCAGACAGGAAAAATGAGCGGC | GAATGACGAAAACGCGCTTGTC | 506 |
| INK1328 | EG6180 *air-1* R62A | CGAGAAGAAAACGAGACGAA | ctggagtctcgacgatttcgacgttggcgctccacttggaaagggaaagtttggaaatgtgttcatcagtgccgagaagaaaacgagacgaattatcgctctgaaagttctcttcaaaacgc | TCAGACAGGAAAAATGAGCGGC | GAATGACGAAAACGCGCTTGTC | 506 |
| INK1376 | *klmt-1* FARS-3-like helix I | AAATGTGGCTGCGAATGCAT | ggactttcgtttcctagatgcaaagtcgattgagaagataaatgtggctcgcgagactgcgcaggtcttccacccatatttatccggtgcgattgttcgccagatttcattag | TGGACAAGAAGCACGGAGAGAC | GGCGGTATCATCACTTCGAGGA | 920 |
| INK1396 | *klmt-1::GFP/mScarlet-spd-5* | CCAGTGTTTTTTCAATATGG | ttctccgttttttcttcaatcaattcacgttttccagtgttttttcaatatggactcaacagaggccgtcatcaaggagttcatgcgtttcaaggtccacatggagggatccatgaacggacacgagttcgagatcgagggagagggagagggacgtccatacgagggaacccaaaccgccaagctcaaggtcaccaaggtaagtttaaacatatatatactaactaaccctgattatttaaattttcagggaggaccactcccattctcctgggacatcctctccccacaattcatgtacggatcccgtgccttcatcaagcacccagccgacatcccagactactggaagcaatccttcccagagggattcaagtgggagcgtgtcatgatcttcgaggacggaggaaccgtctccgtcacccaagacacctccctcgaggacggaaccctcatctacaaggtcaagctccgtggaggaaacttcccaccagacggaccagtcatgcaaaagcgtaccatgggatgggaggcctccaccgagcgtctctacccagaggacgtcgtcctcaaggtaagtttaaacagttcggtactaactaaccatacatatttaaattttcagggagacatcaagatggccctccgtctcaaggacggaggacgttacctcgccgacttcaagaccacctacaaggccaagaagccagtccaaatgccaggagccttcaacatcgaccgtaagctcgacatcacctcccacaacgaggactacaccgtcgtcgagcaatacgagcgttccgtcgcccgtcactccaccggaggatcaggaggatcaggtggttctatggaggataattcagtgttgatggaagactcaaatctcgaagaagtc | GCTTCGAATTCAAAAATAGTGGGCG | CTTCAACTCCTCGTTCTCGTGC | 1279 |
| INK1527 | *fars-3* E292A | GAGGTAGAATACGAGGAGAC | ctgccaaaaaccgttccatgtcgaacaggtggaggtagaatatgctgagaccggagaaaaggagctctatccgcttctctcctatcgagaa | AACCAGACGAAGGAGTACACGG | GCGTGCAGAATATCGTGTCGAG | 524 |
| INK1530 | *fars-3* E292K | GAGGTAGAATACGAGGAGAC | ctgccaaaaaccgttccatgtcgaacaggtggaggtagaatataaagagaccggagaaaaggagctctatccgcttctctcctatcgagaa | AACCAGACGAAGGAGTACACGG | GCGTGCAGAATATCGTGTCGAG | 524 |
| INK1533 | *klmt-1* E313K | GATTGAATACGAGAAATTCG | gaaaaaccacttgtcgttgagcccgttttgattgaatacgagaagtttaaggaaccgagagccttagagctctccccaccgctctcttatcgc | ACGTTGTACTCATCTGACAGCCA | TTTTTGTAGGCACACGCGACTC | 539 |
| INK1574 | *klmt-1* E310K | GATTGAATACGAGAAATTCG | cgaattgaaaaaccacttgtcgttgagcccgttttgattgaatacaaaaagttcgaggaaccgagagccttagagctctccccaccgctctcttatcg | ACGTTGTACTCATCTGACAGCCA | TTTTTGTAGGCACACGCGACTC | 539 |
| INK1601 | KLMT-1 NIL *air-1* A62R | GAAATGTGTTCATTTCCCGCG | gttggccggccattgggaaagggaaagtttggaaatgtgttcatttcccgcgagaagaaaacgagacgaattatcgctctgaaagttctcttcaaaacg | TCAGACAGGAAAAATGAGCGGC | GAATGACGAAAACGCGCTTGTC | 506 |
| INK1690 | *klmt-1* FARS-3-like helix II | AAATGTGGCTGCGAATGCAT | ggactttcgtttcctagatgcaaagtcgattgagaagataaatgtggctcgcgagactgcgcaattccacccatatttatccggtgcgattgttcgccagatttcattag | TGGACAAGAAGCACGGAGAGAC | GGCGGTATCATCACTTCGAGGA | 920 |
| iNK1717 | *klmt-1::GFP ∆N* | CTGATTTTCGGACATTTCTC,  CCTTCCTCCAACTACTCCAG | gttttttccgtggtttttctttgctattttcgaatttaaattaaaaatacggctaataatctcaatattccagagaaatgactccagtggtacgccattacgtttcttttattttttttatattaaaaaataacaatttaaacgacttttcgccagaaaaaccacattttcg | ATCGCATGCGCCTTAAATACCG | AGGAGAATGCAGTTTTTCAGCGG | 459 |
| INK1770 | *klmt-1::GFP ∆C* | CTCCTCGTCACAGAATCCAA,  CTCTGAATCATTAACAAATTC | ctgaagaaagtgactgccaaagtgctcaccctaatgaaaggagaagtcaaaagataaatacattctttcccccctttccccgcttcatcaatgtttcgaag | ATACGGATTTCAGGAGCCACCAA | AAAATCTGGAGTTTACGCGCGTT | 395 |

Table S4.

Yeast strain and plasmids used for complementation assays.

| Strain name | Species | Genotype | Source |
| --- | --- | --- | --- |
| YFL022C  (FRS2) | *S. cerevisiae* | *MATa/α ura3Δ0/ura3Δ0 leu2Δ0 his3Δ1/his3Δ1 lys2Δ0/LYS2 met15Δ0/MET15 can1Δ::LEU2MFA1pr::His3/CAN1 yfgΔ::KanMX/YFG* | Ben-Aroya et al., 2008 |

| Plasmid ID | Name | Description | Use |
| --- | --- | --- | --- |
| p42Nat | p42Nat | Multicopy with cloNat resistance marker for single Gateway insertion | empty vector |
| pCEV-G2-Km | pCEV-G2-Km | TEF-1 and PGK-1 promoter controlled expression cassettes with G418 resistance for yeast transformation | empty vector |
| pAB0165 | pCEV-G2-KM::fars1::fars3 | Constitutive expression of *C.tropicalis* fars-1 (pPGK-1) and fars-3 (pTEF-1) | FARS-1/FARS-3 for complementation |
| pAB0166 | p42Nat::pGal1::KLMT-1 | Galactose inducible expression of KLMT-1; pGal-1 amplified from p406Gal1 | inducible KLMT-1 |

Movie S1-S6. (separate files)

Lineaging and nuclear tracking in control and KLMT-1 expressing embryos.

Data S1. (separate file)

List of genetic crosses performed in this study.

Data S2. (separate file)

Table of heat shock results in wild-type and KLMT-1 expressing *C. elegans* and *C. tropicalis* embryos.

Data S3. (separate file)

Data from tRNA sequencing.

Data S4. (separate file)

Raw data of all quantifications from fluorescence micrographs shown in this study.

Data S5. (separate file)

Descriptions, input sequences and Alphafold3 PTM/iPTM scores for all centrosomal *C. elegans* proteins used in KLMT-1 interaction screening.
